# Inflammation-induced epigenetic memory restores oligodendrocyte progenitor cell regenerative capacity in the aged central nervous system

**DOI:** 10.64898/2026.05.11.724385

**Authors:** Sonia Cabeza-Fernández, Sergio Niñerola, Angela Armengol-Gomis, Juan Paraíso-Luna, Ángeles Casillas-Bajo, Jose A. Gómez-Sánchez, Hugo Cabedo, Ángel Barco, Alerie G de la Fuente

## Abstract

Although remyelination, a central nervous system (CNS) regenerative process mediated by oligodendrocyte progenitor cells (OPCs), takes place in an inflammatory environment the long-term impact of inflammation on OPC remyelination capacity remains unclear. Here, we studied the short- and long-term impact of systemic inflammation on adult OPCs to assess whether transient inflammation triggers enduring chromatin remodelling indicative of inflammatory memory in OPCs. We observed long-lasting epigenetic modifications in response to both lipopolyssaccharide (LPS) and polyinosinic:polycytidylic acid (Poly(I:C)), but only LPS induced a tolerance-like memory. LPS-mediated tolerance-like memory enhanced OPC differentiation after demyelination in aged mice, reducing axonal damage. Our findings reveal OPC epigenetic memory of inflammation as a mechanism by which adult OPCs adapt to inflammatory challenges, which could be harnessed to reduce neuroinflammation and enhance remyelination efficiency in ageing and neurodegenerative diseases.

## INTRODUCTION

The loss or disruption of myelin, essential for nervous system function and axonal health, is a common pathological feature across neurological disorders^1,2^. In the central nervous system (CNS), lost myelin can be restored through remyelination- a regenerative process driven primarily by oligodendrocyte progenitor cells (OPCs)- which differentiate into new myelin-forming oligodendrocytes^1^. Although inflammation is required for efficient remyelination, dysregulated inflammatory responses impair OPC differentiation and hinder repair^3^. Remyelination is highly efficient in young adults but declines with age, contributing to cumulative neurological disability in neurodegenerative disorders such as multiple sclerosis (MS)^4–7^. Understanding the mechanisms that preserve and restore remyelination capacity is therefore a key therapeutic goal, particularly for preventing neurodegeneration in older patients.

Beyond their canonical functions in myelin formation and regeneration, OPCs are increasingly recognised as versatile regulators of CNS function, including immune modulation^8,9^. Inflammatory cues induce immune-related transcriptional programs and OPCs adopt disease-associated states across neurological disorders^10–16^. As CNS-resident progenitors^17–19^, OPCs are likely exposed to diverse inflammatory perturbations throughout life including stress, systemic infection, recurrent inflammatory episodes- such as those occurring in MS relapses-or chronic inflammation characteristic of neurodegenerative diseases and ageing. However, the enduring impact of such exposures on OPC immune and regenerative functions are largely unknown.

Other tissue-resident progenitors, such as epithelial stem cells, adapt to inflammatory cues through epigenetic memory of inflammation (EMI)^20–22^. This form of memory relies on durable epigenetic reprogramming that alters cellular responses to subsequent stressors, thereby shaping their immune-modulatory and regenerative potential^20–23^. OPCs exhibit epigenetic priming of immune gene loci without gene expression in the healthy CNS, late stages of experimental autoimmune encephalomyelitis (EAE) and MS white matter^24–26^. Moreover, neonatal OPCs repeatedly exposed to IFNγ *in vitro* display altered secondary immune responses driven by transcriptomic and epigenetic changes^24^. Together, these observations raise the possibility that OPCs may retain long-lasting molecular imprints of previous inflammatory encounters, acquiring similar forms of EMI that may influence their immune and regenerative functions.

Here we show that adult OPCs acquire a stimulus-specific form of EMI. Acute inflammatory stimulation induces transient transcriptional changes alongside enduring alterations in chromatin accessibility that shape OPC responses to subsequent inflammatory and demyelinating challenges. We find that lipopolysaccharide (LPS), but not polyinosinic:polycytidylic acid (Poly(I:C)), induces a tolerance-like EMI in OPCs that dampens their inflammatory response to following stimuli and enhances OPC differentiation limiting axonal damage during remyelination in aged mice. These findings demonstrate that adult OPCs encode prior inflammatory experiences through stimulus-specific, enduring chromatin remodelling that persists beyond transcriptional resolution. Our data indicate that EMI represents an adaptive feature of adult OPCs capable of modulating aged OPC regenerative potential. Leveraging OPC inflammatory memory could therefore represent a novel therapeutic approach to control neuroinflammation and enhance remyelination during ageing in a range of disease settings.

## RESULTS

### Disease-associated OPC phenotype induced by LPS is transient and coupled to active neuroinflammation

To investigate how OPCs integrate inflammatory signals and define the immediate and long-term consequences of transient inflammation, we used intraperitoneal injection of LPS as a well-established model of acute neuroinflammation^27^. Mice received a single intraperitoneal LPS injection and OPC response was investigated either 24h (LPS 1X) or five weeks (LPS 5w) after stimulation, when systemic inflammation had resolved, and compared to PBS-injected controls (PBS) (**Fig. 1a**). We first investigated whether systemic LPS induced any changes in oligodendrocyte lineage cell density. LPS-driven neuroinflammatory response was accompanied by a transient reduction in oligodendrocyte lineage cell density (**Fig. 1b, c**). This was primarily due to a decrease in the density mature oligodendrocytes 24h after LPS in the ventral white matter of the spinal cord (**Fig. 1b-d**), which was coupled with an increase in OPC density (**Fig. 1e**). However, once inflammation subsided, baseline oligodendrocyte lineage cell density was restored (**Fig. 1b-e**). Despite the transient depletion of mature oligodendrocytes neither the total number of axons nor the density of myelin-coated axons were affected (**Fig. 1f-h**).

**Figure 1:**
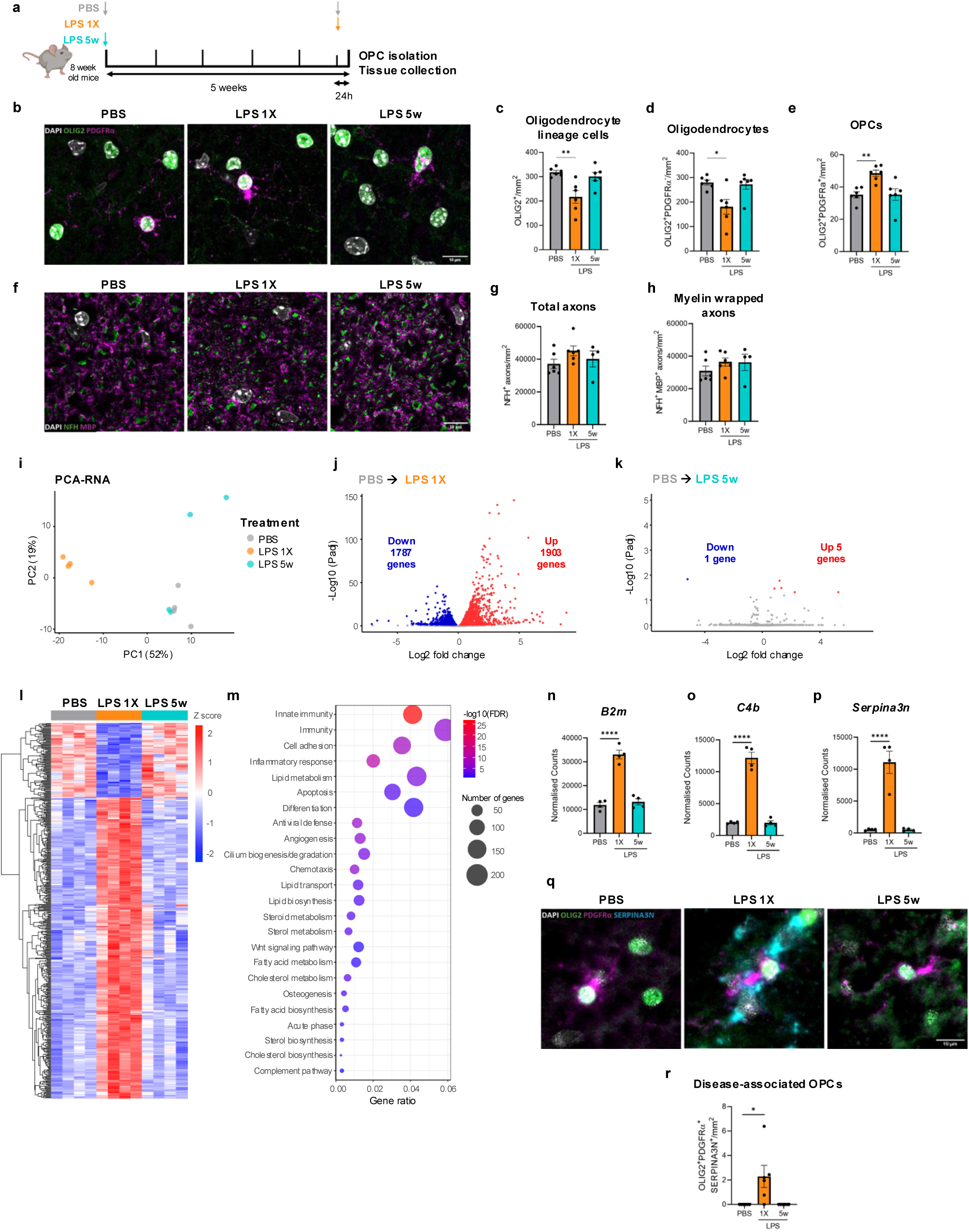
LPS induces a transient inflammatory response in OPCs that returns to baseline at transcriptional level once inflammation is resolved. **a.** Experimental design diagram: Mice were intraperitoneally injected with PBS or LPS and sacrificed either 24h (LPS 1X) or 5 weeks (LPS 5w) later for OPC isolation and tissue collection. **b.** Representative images showing OPCs and mature oligodendrocytes in the spinal cord white matter from mice treated with PBS, LPS 1X and LPS 5w labelled with DAPI (white), OLIG2 (green) and PDGFRα (magenta). Scale bar=10µm. **c-e**. Quantification of the density of total oligodendrocyte lineage cells (OLIG2^+^) **(c)**, mature oligodendrocytes (OLIG2^+^PDGFRα^-^) **(d)**, and OPCs (OLIG2^+^PDGFRα^+^) **(e)** in the spinal cord white matter in PBS, LPS 1X and LPS 5w mice (n=6, 1-way ANOVA, Dunnet’s posthoc test). **f.** Representative immunofluorescence images showing the density of axons and myelin-coated axons in the spinal cord white matter from mice treated with PBS, LPS 1X and LPS 5w labelled with NFH (green) and MBP (magenta). Scale bar =10µm. **g, h.** Quantification of the density of total axons (NFH^+^) **(g)** and myelin wrapped axons (NFH^+,^ MBP^+^) **(h)**, in the spinal cord white matter in PBS, LPS 1X and LPS 5w mice. (n=6, 1-way ANOVA, Dunnet’s posthoc test). **i.** PCA plot representing the clustering of OPCs in PBS, LPS 1X and LPS 5w groups according to their transcriptome. **j.** Volcano plot showing differentially expressed genes (DEGs) in OPCs 24 hours after LPS injection compared with PBS controls. Genes upregulated are shown in red while genes downregulated are depicted in blue. **k.** Volcano plot showing DEGs in OPCs 5 weeks after LPS injection compared with PBS controls. Genes upregulated are shown in red while genes downregulated are depicted in blue. **l.** Heatmap indicating Z score expression levels of the top 500 DEGs in OPCs 24h after LPS in all three groups: PBS, LPS 1X and LPS 5w. **m.** Bubble plot depicting the GO biological processes significantly enriched among the DEGs in OPCs in LPS 1X compared with PBS. **n-p.** Bar graphs showing the normalised counts in RNA-seq of disease-associated OPC markers in PBS, LPS 1X and LPS 5w groups (n=4, 1-way ANOVA, Dunnet’s posthoc test). **q.** Representative immunofluorescence images of spinal cord ventral white matter of mice treated with PBS, LPS 1X and LPS 5w labelled with DAPI (white), OLIG2 (green), PDGFRα (magenta) and SERPINA3N (cyan); scale bar=10µm. **r**. Bar graphs showing the quantification of OPCs expressing disease associated marker SERPINA3N in PBS, LPS 1X and LPS 5w (mice n=6, 1-way ANOVA, Dunnet posthoc test).

Next, we evaluated the immediate and long-lasting LPS-mediated transcriptomic changes in OPCs by bulk RNA sequencing analysis. PDGFRα^+^ OPCs were isolated by magnetic cell sorting (MACS) from mice in the LPS 1X and LPS 5w groups, and their transcriptome compared with those of OPCs from PBS-treated control mice (**Fig. 1a**). OPCs mounted a rapid transcriptional response to systemic LPS, with 3690 differentially expressed genes (DEGs) detected in the LPS 1X group, including 1903 upregulated and 1787 downregulated genes (**Fig. 1i, j**). By contrast, five weeks after LPS injection, when inflammation had subsided, the OPC transcriptome largely reverted to that of PBS-treated controls, with only six DEGs detected and the majority of LPS 1X-associated DEGs restoring baseline transcription levels (**Fig. 1i, k, l**). As expected, the most strongly regulated DEGs in the LPS 1X condition were enriched for Gene Ontology (GO) biological processes such as inflammatory response, immunity, WNT signalling, cell adhesion, lipid and cholesterol metabolism (**Fig. 1m**).

We next investigated whether systemic LPS exposure also induces a disease-associated OPC transcriptional programme, as observed across different neurological disorders^15^. As described for astrocytes^28^, OPCs upregulated disease-associated markers *B2m*, *C4b* and *Serpina3n* 24h after LPS, which expression returned to baseline by five weeks (**Fig. 1n-p**). Immunohistochemistry for SERPINA3N revealed that, similar to mRNA, SERPINA3N protein expression peaked 24h after LPS in OPCs and disappeared when inflammation subsided (**Fig. 1q, r**). Thus, LPS induces a transient immune-like OPC state coupled to active CNS inflammation.

To determine whether transient inflammation elicits long-lasting cell-intrinsic changes, we isolated OPCs five weeks after LPS and analysed their behaviour *in vitro*, in the absence of any additional exogenous stimulation (**Sup. Fig. 1a**). We observed no differences in migration (**Sup. Fig. 1b**), proliferation (**Sup. Fig. c, d**) or differentiation capacity between OPCs isolated from LPS-treated mice and those from PBS-treated control mice (**Sup. Fig. 1e-g**). Thus, OPCs mount a rapid inflammatory transcriptional response to LPS that returns to baseline once inflammation has resolved.

### LPS induces long-lasting changes in chromatin accessibility in adult OPCs

EMI is characterised by persistent epigenetic remodelling in response to inflammatory cues and has been described across diverse long-lived stem cell populations such as epithelial or intestinal stem cells^20–23^. To determine if adult OPCs also retain enduring chromatin alterations following LPS exposure, we first quantified the number and spatial distribution of histone modification puncta within OPC nuclei in PBS, LPS 1X and LPS 5w groups using super-resolution microscopy, as a proxy for changes in chromatin compartimentalization^29,30^. Lysine 27 acetylation on histone 3 (H3K27ac), a hallmark of active chromatin^31^, exhibited a redistribution towards the centre of the nucleus in both LPS 1X and LPS 5w, without changes in the total number of puncta. (**Fig. 2a-c**). Similarly, lysine 27 tri-methylation on histone 3 (H3K27me3)-a hallmark of repressed chromatin domains^31^- showed a modest but significant redistribution towards the nuclear periphery in LPS 1X OPCs, again without changes in overall H3K27me3 puncta number. In contrast, OPCs from the LPS 5w group displayed an increased number of H3K27me3 puncta while maintaining the nuclear distribution comparable to that of control OPCs (**Fig. 2d-f**).

**Figure 2:**
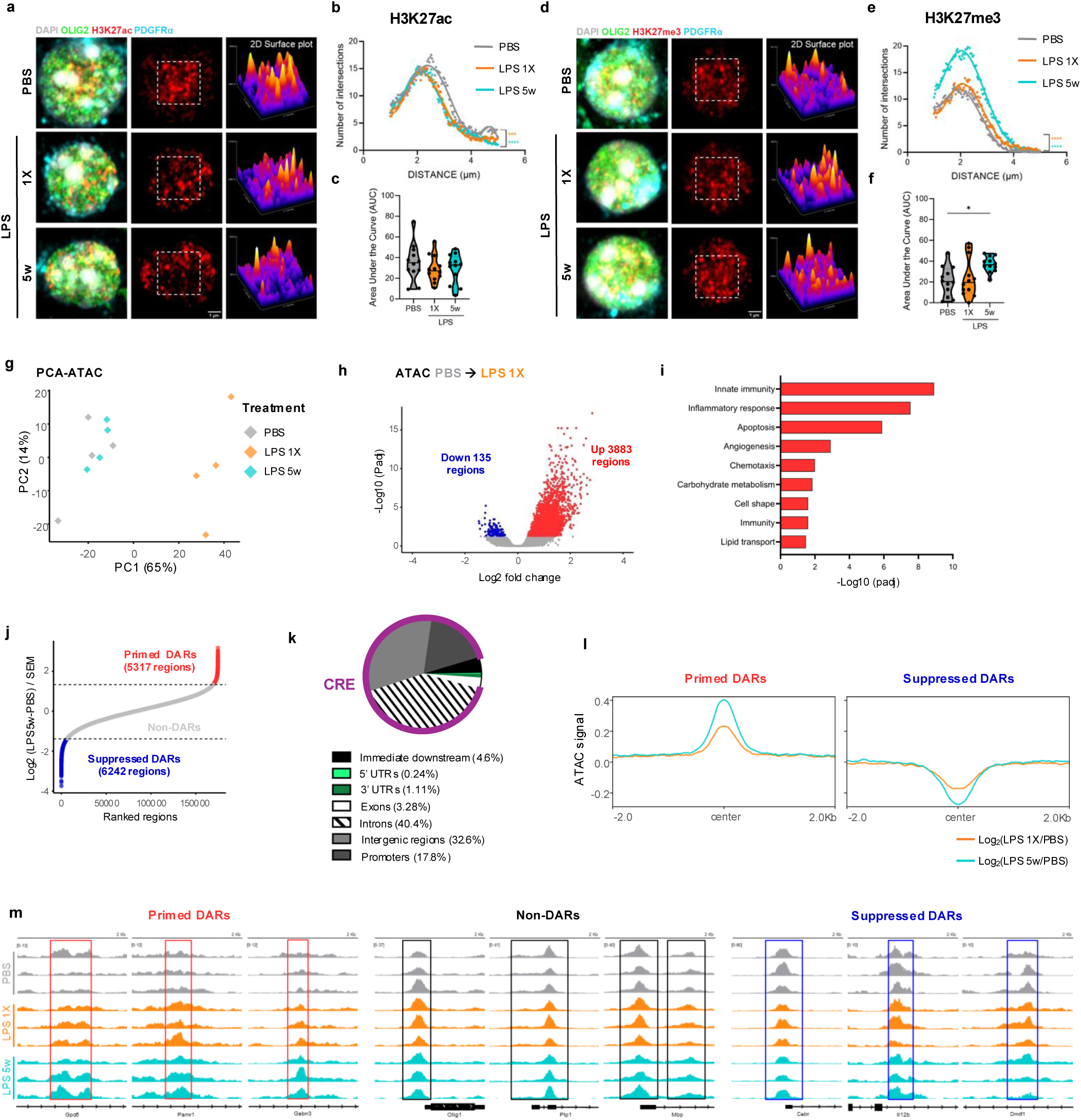
LPS induces long-lasting changes in chromatin accessibility in OPCs. **a.** Super-resolution images showing the organization of H3K27ac domains in the nuclei of OPCs in the spinal cord white matter in PBS, LPS 1X and LPS 5w labelled with DAPI (white), OLIG2 (green), H3K27ac (red), PDGFRα (cyan) and a 2D surface plot representation of the H3K27ac distribution and intensity in the inset; scale bar=1µm. **b**. H3K27ac puncta radial distribution analysis in OPC nuclei in mice treated with PBS, LPS 1X and LPS 5w measured using Sholl analysis (2-way ANOVA, Tukey’s multiple comparison test, n=12 cells, n=6 mice). **c.** Changes in the total number of H3K27ac puncta measured as area under the curve analysis (1-way ANOVA, Dunnet’s posthoc test, n=12 cells, n=6 mice). **d.** Super-resolution images showing the organization of H3K27me3 puncta in the nuclei of OPCs in the spinal cord white matter in PBS, LPS 1X and LPS 5w labelled with DAPI (white), OLIG2 (green), H3K27me3 (red), PDGFRα (cyan) and a 2D surface plot representation of the H3K27me3 distribution and intensity in the inset; scale bar=1µm. **e**. H3K27me3 puncta radial distribution analysis in OPC nuclei in mice treated with PBS, LPS 1X and LPS 5w measured using Sholl analysis (2-way ANOVA, Tukey’s multiple comparison test, n=12 cells, n=6 mice). **f.** Changes in the total number of H3K27me3 domains determined by area under the curve analysis (1-way ANOVA, Dunnet’s posthoc test, n=12 cells, n=6 mice). **g.** PCA plot depicting the clustering of OPCs in PBS, LPS 1X and LPS 5w groups based in chromatin accessibility. **h.** Volcano plot comparing chromatin accessibility between PBS and LPS 1X. **i.** Bar graph showing GO biological processes enriched in upregulated DARs in LPS 1X when compared to PBS. **j.** Elbow plot comparing PBS and LPS 5w chromatin accessibility ranked by the Wald statistic divided by the s.e.m. Primed DARs (red) are identified as regions that are more accessible in LPS 5w than PBS, while suppressed DARs (blue) represent chromatin regions that are less accessible in LPS 5w than in PBS control. **k.** Pie chart indicating the type of DNA region associated with memory DARs (primed and suppressed) identified in the Elbow plot comparing LPS 5w and PBS. **l.** Chromatin accessibility profile representing log_2_ fold change in chromatin accessibility for primed and suppressed DARs in PBS, LPS 1X and LPS 5w measured as distance from the peak center. **m.** Representative Integrative Genomic View (IGV) images of the ATAC signal profile of three individual replicates showing examples of genomic loci of interest classified as primed, unchanged and suppressed regions in PBS, LPS 1X and LPS 5w OPCs. Boxes highlight ATAC peaks. Genes associated with these chromatin regions are as indicated.

To establish whether this enduring changes in histone modification distribution observed in OPC nuclei reflect changes in chromatin accessibility, we performed bulk assay for transposase accessible chromatin sequencing (ATAC-seq) on OPCs isolated from PBS-, LPS 1X- and LPS 5w-treated mice. As expected, the largest changes in chromatin accessibility occurred immediately after LPS exposure, with 4018 differentially accessible regions (DARs) identified in the LPS 1X group, the majority of which (3883) exhibited increased accessibility relative to OPCs isolated from PBS-treated controls (**Fig. 2g, h**). These DARs significantly correlated with the transcriptional changes observed in LPS 1X (**Sup. Fig. 2a**) and were also enriched in GO terms associated with inflammation, innate immunity, angiogenesis and lipid transport (**Fig. 2i**), consistent with a rapid OPC inflammatory response.

By five weeks post-LPS injection, once inflammation had subsided, global chromatin had largely returned to baseline levels (**Fig. 2g**). However, despite this apparent normalization, comparison of log_2_ fold-change between PBS- and LPS 5w- derived RNA-seq and ATAC-seq data revealed no significant correlation, indicating a partial uncoupling between chromatin accessibility and transcription at this time point (**Sup. Fig. 2b**). To capture the chromatin feature underlying this uncoupling and corresponding to the persistent differences in histone modification distribution observed between PBS and LPS 5w OPCs (**Fig. 2a-f)**, we next compared chromatin accessibility profiles between these two groups^22,32,33^.

This analysis identified 11559 memory DARs, including 5317 DARs with increased accessibility in LPS 5w OPCs relative to PBS controls (primed DARs) and 6242 DARs with decreased accessibility (suppressed DARs), while approximately 93% of the chromatin regions remained unchanged (non-DARs) (**Fig. 2j**). Most memory DARs mapped to intronic (40.4%) or intergenic regions (32.6%), with only 17.8% assigned to promoters (**Fig 2k**), indicating preferential localisation to cis-regulatory elements (CRE) such as enhancers, consistent with findings in other stem and glial cell populations^21,22,32^. Both primed and suppressed DARs displayed only subtle accessibility changes between PBS and LPS 1X group, with the highest change in accessibility emerging at the LPS 5w timepoint (**Fig. 2l, m**; **Sup. Fig. 2c, d**). GO term biological process analysis of the genes annotated to primed DARs by proximity revealed enrichment for processes related to cell migration, cell adhesion, ion signalling or lipid transport (**Sup. Fig. 2e**), whereas suppressed DARs were enriched for terms associated with nervous system development, synaptic membrane, transcription, or cell differentiation (**Sup. Fig. 2f**). Together, these results indicate that despite global recovery in chromatin accessibility, a discrete subset of regulatory elements acquire long-lasting accessibility changes in adult OPCs, suggesting persistent chromatin accessibility changes in loci that may modulate OPC response to future challenges, including OPC regenerative response.

### LPS pre-exposure induces a tolerance-like memory response in adult OPCs

EMI represents an adaptative response to inflammatory cues that can manifest as either training, an enhanced secondary inflammatory response that promotes pathogen clearance or accelerates repair, or tolerance, in which repeated stimulation elicits an attenuated inflammatory response to limit tissue damage^34–36^. To determine whether the persistent changes in chromatin accessibility observed in OPCs reflect EMI, we isolated OPCs 24h after either a single LPS injection (LPS 1X) or the second of two LPS injections separated by five weeks (LPS 2X) and compared their transcriptomic and chromatin accessibility profiles (**Fig. 3a**). Bulk RNA sequencing revealed that OPCs from the LPS 2X group clustered in between PBS and LPS 1X samples (**Fig. 3b**). Direct comparison of the transcriptomic differences between LPS 1X and LPS 2X OPCs identified 941 DEGs (**Fig. 3c**), of which 394 genes were downregulated and enriched in GO terms such as inflammation, innate immunity, chemotaxis and cell adhesion (**Sup. Fig. 3a**), while 547 genes were upregulated and enriched for cilium biogenesis, cell differentiation, and cell adhesion (**Sup. Fig. 3b**), consistent with an attenuated inflammatory response. Although the log_2_ fold change of all genes showed a strong positive correlation between LPS 1X vs PBS and LPS 2X vs PBS comparisons (**Sup. Fig. 3c**), the top 500 DEGs identified between PBS and LPS1X comparison exhibited a markedly reduced expression changes in OPCs isolated from the LPS 2X group (**Fig. 3d**). This blunted inflammatory response was further supported by the direct comparison of OPCs from PBS and LPS 2X conditions, which identified only 329 DEGs (**Sup. Fig. 3d**). Of these, approximately 89% of the downregulated genes and 94% of the upregulated genes overlapped with LPS 1X DEGs (**Sup. Fig. 3e, f**). This blunted inflammatory response was replicated in a second cohort using qPCR, where inflammatory and disease-associated genes such as *Tnf*, *C4b* and *Serpina3n* were robustly induced in LPS 1X OPCs, but not in LPS 2X OPCs (**Fig. 3e-g**). Thus, OPCs mount a transcriptional response of lower magnitude upon repeated LPS exposure, consistent with tolerance-like inflammatory memory.

**Figure 3:**
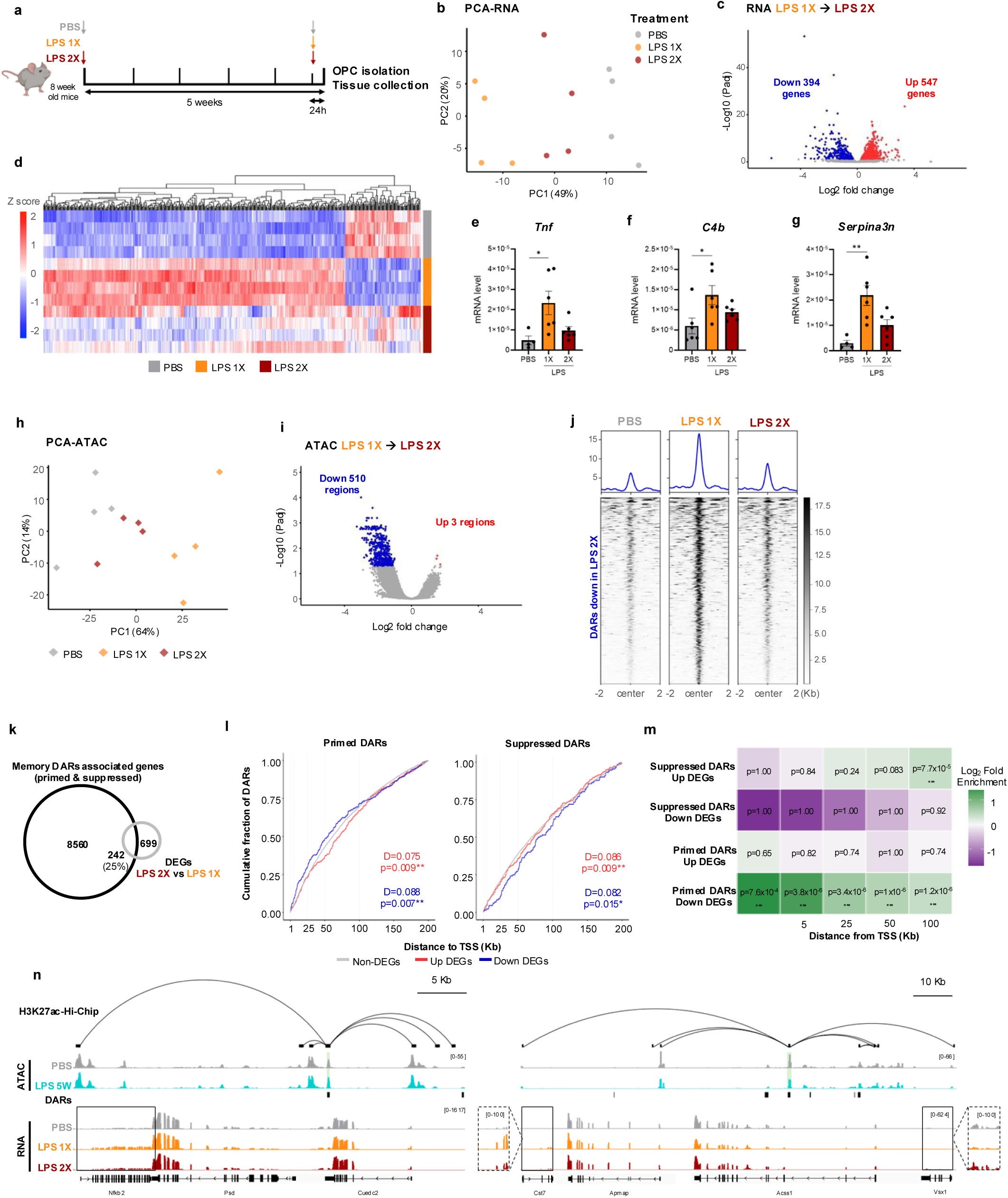
LPS pre-exposure induces a tolerance-like OPC response to a second LPS stimulus. **a.** Schematic diagram depicting the experimental design: Mice were exposed to PBS, a single LPS injection (LPS 1X) or repeated LPS injections separated in time (LPS 2X) and sacrificed for OPC isolation and tissue collection 24h after the last injection. **b.** PCA plot showing the clustering of OPCs in PBS, LPS 1X and LPS 2X groups. **c.** Volcano plot depicting the DEGs identified in LPS 2X mice when compared to LPS 1X (upregulated in red, downregulated in blue). **d.** Heatmap displaying the Z score changes in expression of the top 500 DEGs induced in LPS 1X in the repeated LPS 2X group. **e-g.** mRNA expression level changes for inflammatory (*Tnf*) (**e**) and disease-associated markers (*C4b* (**f**), *Serpina3n* (**g**)) measured by qPCR in OPC isolated after PBS, LPS 1X and LPS 2X treatment (n=4, 1-way ANOVA, Dunnett’s posthoc test). **h.** PCA plot showing the clustering of PBS, LPS 1X and LPS 2X OPCs according to chromatin accessibility. **i.** Volcano plot showing DARs between LPS 1X and LPS 2X OPCs. DARs with increased chromatin accessibility are shown in red while DARs with diminished chromatin accessibility are depicted in blue. **j.** Chromatin accessibility profile and heatmap showing the mean ATAC signal of the down DARs identified between LPS 2X and LPS 1X in OPCs from PBS, LPS 1X and LPS 2X. **k.** Venn diagram showing common genes between memory DARs and DEGs between LPS 2x and 1X. **l.** Empirical cumulative distribution function (ECDF) plots showing the cumulative fraction of genes as a function of their distance to the nearest primed or suppressed DAR. Curves for DEGs and non-significant genes are compared visually and tested using the Kolmogorov–Smirnov test. **m.** Heatmap showing the fold enrichment and significance of log_2_ fold-enrichment of DAR–DEG pairings across TSS-centered windows (≤100 kb) (Hypergeometric test, Benjamini-Hochberg correction). **n**. Representative examples of H3K27ac Hi-Chip contacts formed by primed and suppressed DARs identified between PBS and LPS 5w (marked with black and green shadow) with DEGs identified between LPS 1X and LPS 2X (outlined with black box). Merged ATAC-seq and RNA-seq profiles from four independent samples in PBS, LPS 5w, LPS 1X and LPS 2X conditions are shown. Zoom-in panels display RNAseq profiles at the regions marked by rectangles.

Next, we examined whether tolerance-like EMI was also evident at the level of chromatin accessibility. Consistent with the transcriptomic data, OPCs isolated from LPS 2X group also clustered between PBS and LPS 1X treated samples based on chromatin accessibility changes (**Fig. 3h**). Direct comparison between LPS 1X and LPS 2X OPCs identified 513 DARs, with a predominant loss of chromatin accessibility in the LPS 2X condition (**Fig. 3i**). Most of these DARs corresponded to loci that gained accessibility after a single LPS injection but failed to reopen following the second LPS stimulus (**Fig. 3j).** Overall, LPS 2X OPCs exhibited an attenuated chromatin response, with only 8 DARs detected between PBS and LPS 2X conditions and the majority of the DARs identified in the PBS and LPS 1X comparison no longer differentially accessible (**Sup. Fig. 3g-k**).

To determine if the persistent chromatin remodelling observed in the LPS 5w condition was associated with the tolerance-like EMI response detected after repeated LPS exposure, we investigated whether primed and suppressed DARs might regulate transcriptional changes observed between LPS 1X and LPS 2X OPCs. We first quantified the proportion of DEGs between LPS 1X and LPS 2X OPCs that were associated with a nearby primed or suppressed DAR and found that approximately 25% of these DEGs have a memory DAR annotated by proximity (**Fig. 3k**). We next analysed whether memory DARs exhibited preferential spatial coupling with DEGs. DEGs were consistently located significantly closer to memory DARs than non-DEGs, whereas no difference in proximity between DEGs and non-DEGs was detected for regions without differential accessibility (**Fig. 3l, Sup. Fig. 3l**). Primed DARs were enriched within short-range transcription start site (TSS)-centered windows (±1-5kb), while suppressed DARs were depleted from these regions and instead preferentially associated with upregulated DEGs at more distal enhancer-like ranges (∼100kb) (**Fig. 3m**). Permutation based modelling confirmed that these proximity patterns were unlikely attributable to chance (**Sup. Fig. 3m**). To examine whether distal memory DARs could regulate DEGs through higher order 3D chromatin organisation, we mapped their long-range 3D genomic interactions using publicly available H3K27ac Hi-ChIP data from control OPCs^25^. Several memory DARs identified between PBS and LPS 5w engaged in long-range contacts with DEGs identified between LPS 1X and LPS 2X OPCs (**Fig. 3n**). Together, these data combined with the increased nuclear abundance of the repressive histone mark H3K27me3 in LPS 5w OPCs (**Fig. 2d-f**), suggest that memory DARs may directly contribute to the attenuated secondary transcriptional and chromatin accessibility responses observed following repeated LPS exposure, thereby driving a tolerance-like form of EMI in adult OPCs.

### Epigenetic inflammatory memory in OPCs is stimulus-specific

In innate immune and stem cells EMI can provide cross-protection independently of the initiating stimulus^20,23,34,36,37^. However, the magnitude and functional impact of this memory depend on the nature, dose and duration of the inflammatory exposure^23,34–38^. To determine whether adult OPCs harbour EMI independent of the inflammatory trigger or instead establish distinct EMI programmes in response to different inflammatory cues, we repeated our experimental paradigm using polyinosinic:polycytidylic acid (Poly(I:C)), a double stranded RNA that elicits antiviral responses and induces mild CNS inflammation^39^. OPCs mounted a rapid transcriptomic response 24h after Poly(I:C) administration (Poly(I:C) 1X), which returned to baseline by five weeks (Poly(I:C) 5w) (**Fig. 4a**). We identified 3351 DEGs between PBS and Poly(I:C) 1X OPCs (**Fig. 4b**), enriched for GO terms related to inflammatory responses, antiviral defense and lipid metabolism, indicating a partial overlap with the LPS-induced response (**Sup. Fig. 4a**). Consistently, approximately 56% of Poly(I:C) 1X DEGs overlapped with those induced by LPS 1X (**Sup. Fig. 4b**). By contrast, only six DEGs were detected between PBS and Poly(I:C) 5w OPCs, none of which overlapped with DEGs identified in LPS 5w OPCs (**Fig. 4c**, **Sup. Fig 4c**). In line with previous reports^40^, Poly(I:C) transiently induced disease-associated marker expression (*B2m*, *C4b*, *Serpina3n*), which returned to baseline by five weeks, supporting the notion that disease-associated transcriptional states are coupled to active CNS inflammation (**Sup. Fig. 4d-f**). Despite showing a similar trend towards reduced oligodendrocyte lineage cell and mature oligodendrocyte density, Poly(I:C), unlike LPS, did not significantly alter oligodendrocyte lineage cell density nor total or myelin-coated axonal density at either 24h or five weeks (**Sup. Fig. 4g-k**).

**Figure 4:**
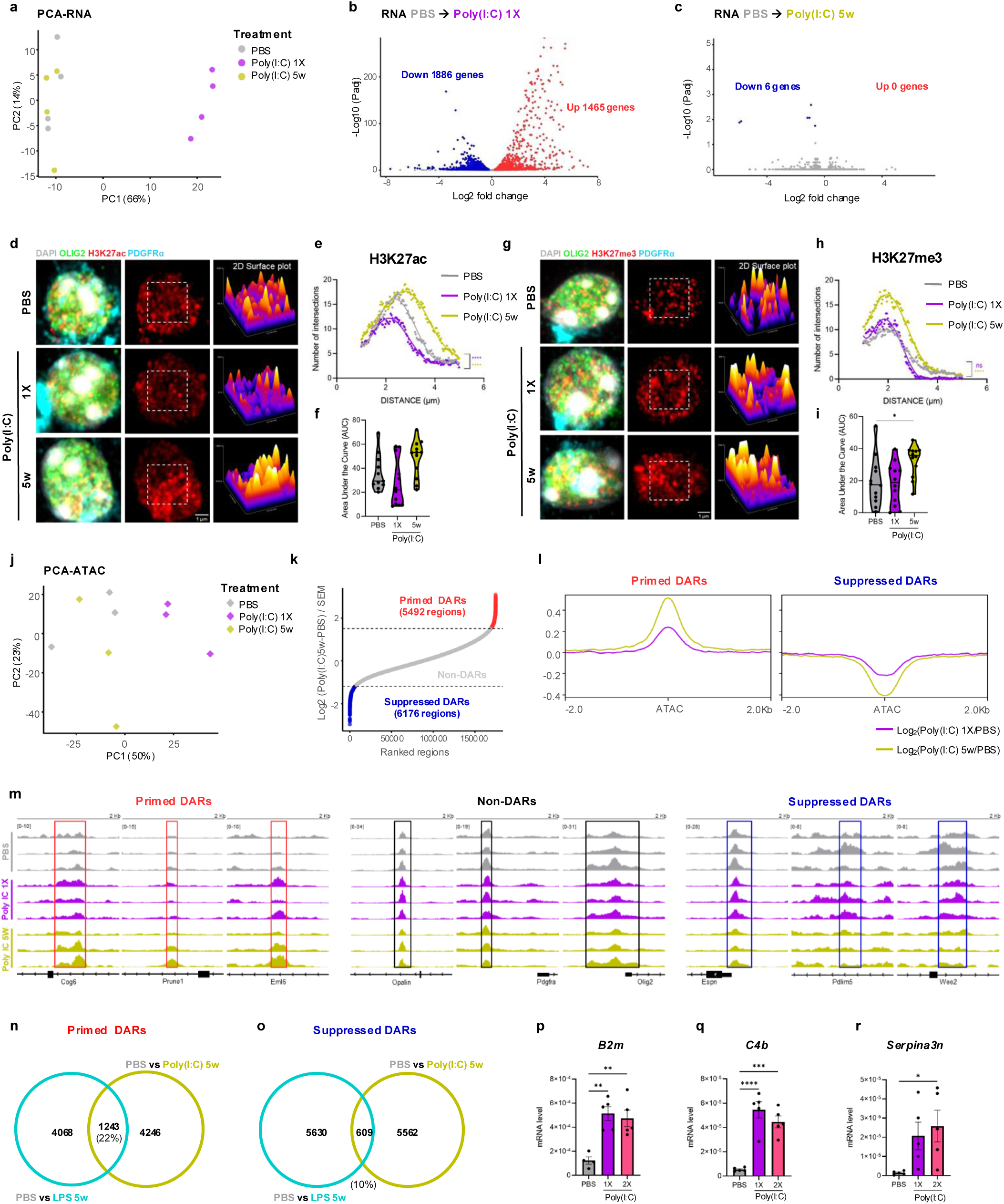
Poly(I:C) also induces a transient OPC response at transcriptomic level and long-lasting epigenetic reprogramming but different to that of LPS. **a.** PCA plot showing the clustering of OPCs isolated from PBS, Poly(I:C) 1X and Poly(I:C) 5w based on their transcriptome. **b, c.** Volcano plot showing DEGs in OPCs 24 hours (**b**) and 5-weeks (**c**) after Poly(I:C) injection compared with PBS controls. Genes upregulated are shown in red while genes downregulated are depicted in blue. n=4. **d.** Super-resolution images showing the organization of H3K27ac domains in the nuclei of OPCs in the spinal cord white matter in PBS, Poly(I:C) 1X and Poly(I:C) 5w labelled with DAPI (white), OLIG2 (green), H3K27ac (red), PDGFRα (cyan) and a 2D surface plot representation of the H3K27ac distribution and intensity in the inset; scale bar=1µm. **e**. H3K27ac puncta radial distribution analysis in OPC nuclei in mice treated with PBS, Poly(I:C) 1X and Poly(I:C) 5w measured using Sholl analysis (2-way ANOVA, Tukey’s multiple comparison test, n=11-12 cells, n=6 mice). **f.** Changes in the total number of H3K27ac puncta determined by area under the curve analysis comparing PBS, Poly(I:C) 1X and Poly(I:C) 5w (1-way ANOVA, Dunnet’s posthoc test, n=11-12 cells, n=6 mice). **g.** Super-resolution images showing the organization of H3K27me3 puncta in the nuclei of OPCs in the spinal cord white matter in PBS, Poly(I:C) 1X and Poly(I:C) 5w labelled with DAPI (white), OLIG2 (green), H3K27me3 (red), PDGFRα (cyan) and a 2D surface plot representation of the H3K27me3 distribution and intensity in the inset; scale bar=1µm. **h**. H3K27me3 puncta radial distribution analysis in OPC nuclei in mice treated with PBS, Poly(I:C) 1X and Poly(I:C) 5w measured using Sholl analysis (2-way ANOVA, Tukey’s multiple comparison test, n=12 cells, n=6 mice). **i.** Changes in the total number of H3K27me3 puncta determined by area under the curve analysis comparing PBS, Poly(I:C) 1X and Poly(I:C) 5w (1-way ANOVA, Dunnet’s posthoc test, n=12 cells, n=6 mice). **j.** PCA plot showing the clustering of OPCs isolated from PBS, Poly IC 1X and Poly(I:C) 5w based on their chromatin accessibility. **k.** Elbow plot comparing PBS and Poly(I:C) 5w chromatin accessibility ranked by the Wald statistic divided by the s.e.m. Primed domains (red) are identified as domains that are more accessible in Poly(I:C) 5w than PBS, while suppressed domains (blue) represent chromatin regions that are less accessible in Poly(I:C) 5w than in PBS control. **l.** Chromatin accessibility profile of log_2_ change in chromatin accessibility (ATAC signal) for primed and suppressed domains in Poly(I:C) 1X and Poly(I:C) 5w. **m.** Representative IGV profile images of the ATAC signal of three individual replicates of example genomic loci classified as primed, unchanged and suppressed domains in PBS, Poly(I:C) 1X and Poly(I:C) 5w OPCs are shown. Genes associated with these chromatin regions are as indicated. **n, o.** Venn diagrams showing the primed (**n**) and suppressed (**o**) memory domains shared between LPS 5w and Poly IC 5w when compared with PBS. **p-r.** mRNA expression level of disease-associated OPC markers in PBS, Poly(I:C) 1X and Poly(I:C) 2X groups measured by qPCR (n=4-5, 1-way ANOVA, Dunnet’s posthoc test).

We next determined whether Poly(I:C) also induced long-lasting epigenetic changes in OPCs. As a first approximation, we analysed the spatial distribution and number of H3K27ac and H3K27me3 histone modifications in OPC nuclei, as described above. OPCs from Poly(I:C) 1X-treated mice exhibited a pronounced re distribution of H3K27ac puncta towards the nuclear center whereas at five weeks (Poly(I:C) 5w) H3K27ac redistributed towards the nuclear periphery, without changes in overall puncta number (**Fig. 4d-f**). In contrast, H3K27me3 puncta showed no early changes in Poly(I:C) 1X OPCs but increased in number and shifted their distribution toward the nuclear periphery in OPCs from the Poly(I:C) 5w group (**Fig. 4g-i**). To determine whether these changes were associated with persistent changes in chromatin accessibility, we performed bulk ATAC-seq on OPCs isolated from Poly (I:C)-treated mice. As observed following LPS exposure, most Poly(I:C)-induced changes in chromatin accessibility occurred at 24h and were largely resolved by five weeks (**Fig. 4j, Sup. Fig. 4l**). Using the elbow-based approach^22,32,33^ to compare PBS with Poly(I:C) 5w, we identified 5492 primed and 6176 suppressed DARs (**Fig. 4k**). As for LPS, these DARs displayed minimal changes in the Poly(I:C) 1X timepoint, with more pronounced differences emerging between PBS and Poly(I:C) 5w OPCs (**Fig. 4l, m; Sup. Fig. 4m, n**). These majority of Poly(I:C)-induced memory DARs mapped to intronic and intergenic regions, consistent with regulation via enhancer-associated elements (**Sup. Fig. 4o**). Despite these overall similarities in chromatin accessibility dynamics, overlap between memory DARs identified after LPS and Poly(I:C) was limited with only 22% of primed (1243) and 10% of suppressed (609) shared between LPS 5w- and Poly(I:C) 5w (**Fig. 4n, o**), indicating that each inflammatory stimulus establishes a distinct epigenetic signature in OPCs. To determine if Poly(I:C) driven persistent chromatin accessibility changes were also associated with tolerance-like EMI, mice were exposed to two Poly(I:C) injections distanced by five weeks (Poly(I:C) 2X), and OPC gene expression changes evaluated by qPCR. In contrast to LPS, Poly(I:C) 1X and Poly(I:C) 2X induced comparable expression levels of disease-associated markers in OPCs, with no evidence of either tolerance or training (**Fig. 4p-r**). Together, these results suggest that distinct inflammatory stimuli induce qualitatively different long-lasting chromatin accessibility alterations in adult OPCs, and that Poly(I:C), unlike LPS, does not elicit tolerance-like EMI under these conditions.

### LPS pre-exposure induces changes in inflammatory and regenerative pathways and age-related hallmarks in OPCs

EMI not only modifies responses to subsequent exposures to the same insult but can also shape cellular responses to a broad range of subsequent stressors including infections, inflammatory challenges, tissue damage or cancerous events^20,22,23,37^. We therefore asked whether EMI influences the regenerative capacity of OPCs following myelin damage. Because inflammation is required for efficient remyelination^41–43^ but impairs OPC differentiation when chronic^3^, we first assessed whether LPS-induced tolerance-like EMI altered transcriptional programs relevant to remyelination. Using manually curated gene sets associated with regeneration, OPC differentiation and myelin formation we computed Gene Set Variation Analysis (GSVA) enrichment scores across conditions^44^. Consistent with heightened inflammation, OPCs from LPS 1X group exhibited a reduced score in gene signatures associated with regeneration, OPC differentiation and myelination. IN contrast, these signatures were partially restored in OPCs from the LPS 2X group (**Fig. 5a-c**). However, gene signatures associated with stemness, migration or cell growth, as well as pathways specifically linked to myelin regeneration such as lipid metabolism or cholesterol metabolism were unchanged across conditions (**Fig. 5d-h**).

**Figure 5:**
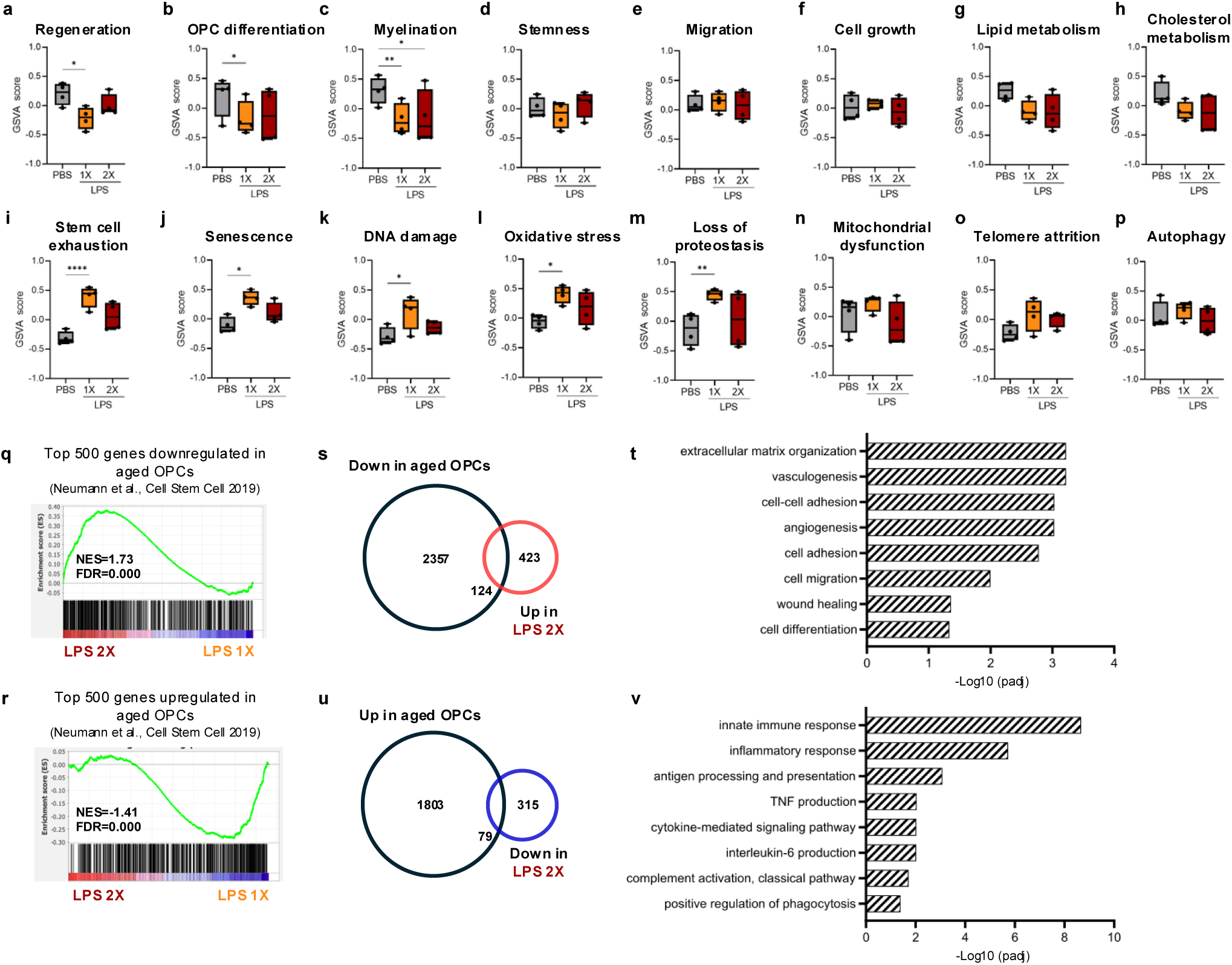
LPS pre-exposure induces changes in inflammatory and regenerative pathways and in age-related hallmarks in OPCs. **a-h.** Box plots depicting changes in gene set variation analysis (GSVA) enrichment scores in manually curated gene signatures associated with regenerative processes and myelin regeneration between PBS, LPS 1X and LPS 2X (n=4, Kruskal Wallis, Tukey’s posthoc test). **i-p.** Box plots depicting changes in GSVA enrichment scores in manually curated gene signatures associated with hallmarks of ageing in PBS, LPS 1X and LPS 2X (n=4, Kruskal Wallis, Tukey’s posthoc test). **q.** Gene set enrichment (GSEA) analysis of the top 500 genes significantly downregulated in aged OPCs and their enrichment in the transcriptome of LPS 2X compared to LPS 1X OPCs. **r.** GSEA analysis of the top 500 genes significantly upregulated in aged OPCs and their enrichment in the transcriptome of LPS 2X compared to LPS 1X OPCs. **s, t.** Venn diagram showing the overlap between the genes significantly upregulated in LPS 2X compared with LPS 1X and the genes significantly downregulated in aged OPCs compared with young OPCs(**s**) and the GO biological processes enriched in the overlapping genes (**t**). **u, v.** Venn diagram showing the overlap between the genes significantly downregulated in LPS 2X compared with LPS 1X and the genes significantly upregulated in aged OPCs compared with young OPCs (**u**) and the GO biological processes enriched in the overlapping genes (**v**).

Genes downregulated in LPS 2X OPCs were predominantly enriched for the GO Terms related to immunity and inflammatory responses (**Sup. Fig. 3a**). As increased inflammation is a hallmark of ageing and a key driver for age-associated functional decline^45^, we next investigated whether LPS-mediated tolerance-like EMI also modulated transcriptional programmes linked to other hallmarks of ageing^45^. GSVA revealed that OPCs from the LPS 1X group displayed increased enrichment score for gene signatures associated with stem cell exhaustion, senescence, DNA damage, oxidative stress and loss of proteostasis. In contrast, these signatures were indistinguishable between OPCs derived from PBS and LPS 2X-treated mice (**Fig. 5i-m**). Other hallmarks of ageing including mitochondrial dysfunction, telomere attrition or autophagy remained unchanged across conditions (**Fig. 5n-p**). To further assess whether LPS-mediated tolerance-like EMI selectively modulated age-related pathways in OPCs, we compared our dataset with a previously published transcriptomic profile of aged OPCs^46^ using Gene Set Enrichment Analysis (GSEA)^47^. The top 500 genes downregulated with age in OPCs were enriched in the LPS 2X group (**Fig. 5q**), while the top 500 genes increased in aged OPCs were enriched in the LPS 1X group (**Fig. 5r**). Among the genes differentially expressed between LPS 1X and LPS 2X OPCs, 124 genes upregulated in LPS 2X condition (22.6%) were significantly downregulated in aged OPCs (**Fig. 5s**). These genes were enriched for GO biological processes including FGF signalling, wound healing, cell migration, cell adhesion, or extracellular matrix, pathways tightly associated to regenerative capacity^48^ (**Fig. 5t**). Conversely, 79 genes downregulated in LPS 2X OPCs (20%) were significantly upregulated in aged OPCs (**Fig. 5u**) and enriched for GO terms related to inflammatory response, cytokine-mediated signalling or antigen processing and presentation (**Fig. 5v**), immune pathways previously associated linked to impaired remyelination^13^. These results suggest that tolerance-like EMI may partially mitigate age-associated transcriptional changes in OPCs.

### LPS-mediated tolerance-like memory enhances OPC regenerative potential with age

Ageing is a major limiting factor for remyelination owing to intrinsic changes in aged OPCs that limit their capacity to respond to damage and differentiate into myelin forming oligodendrocytes, thereby hindering OPC regenerative capacity^4,5,46^. Considering LPS-mediated transcriptional changes observed in OPCs, we next tested whether LPS-induced EMI could help overcome age-associated decline in OPCs regenerative function. Twelve-month-old mice, in which remyelination capacity is already compromised^49–51^, received LPS followed by lysolecithin-induced spinal cord demyelination five weeks later, and tissue was analysed at 7-and 14- days post-lesioning (dpl) (**Fig. 6a**).

**Figure 6:**
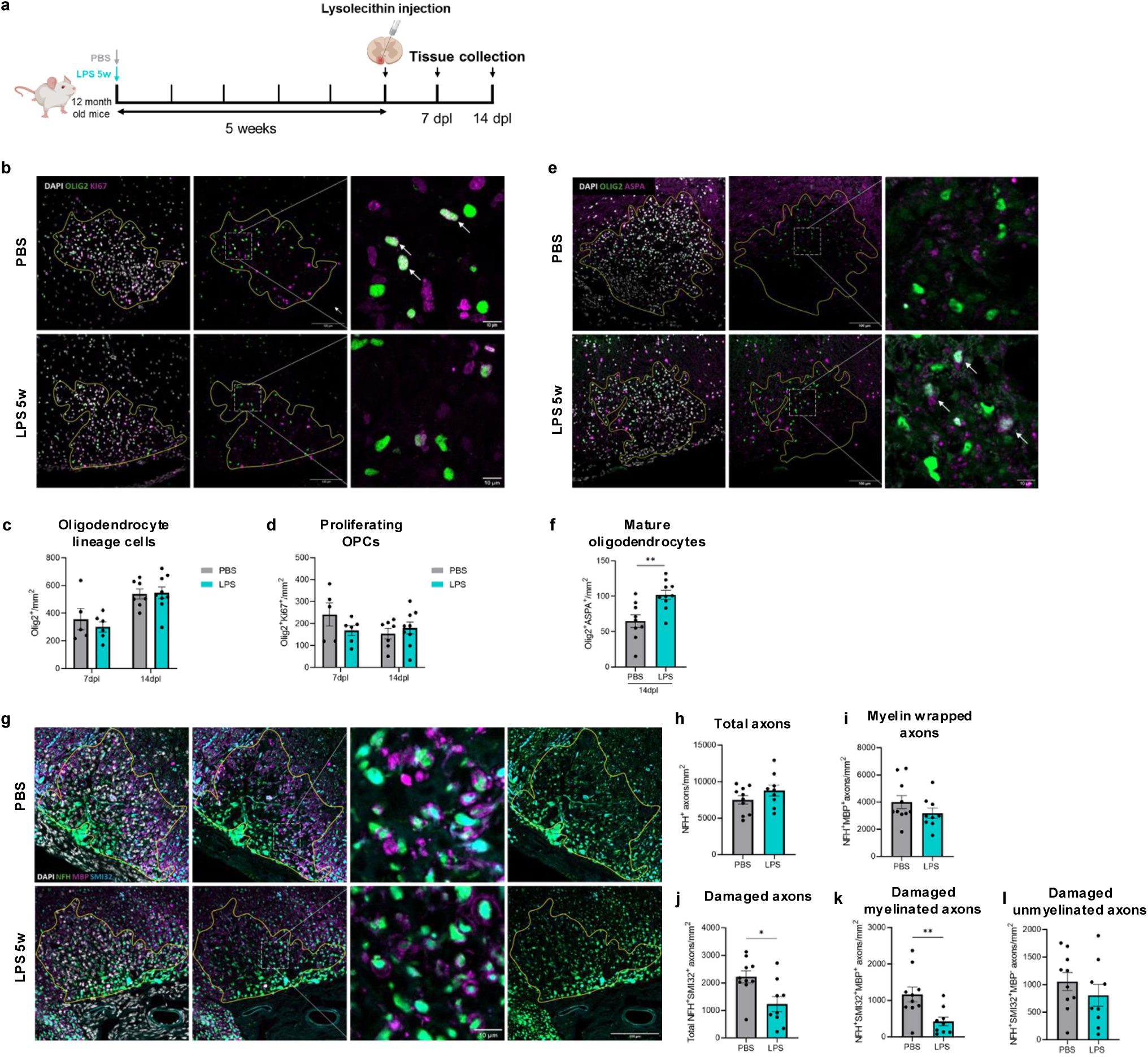
LPS-mediated tolerance-like memory enhances aged OPC regenerative potential. **a.** Schematic diagram representing the experimental approach used to induce OPC inflammatory memory in aged OPCs prior to lysolecithin-induced demyelination. **b.** Representative images of proliferating OPCs labelled with DAPI (white), OLIG2 (green) and KI67 (magenta) in lysolecithin-induced demyelinating lesions in PBS- and LPS-treated mice. Demyelinated area is demarcated by the yellow line. Scale bar=100µm; Inset scale bar=10µm. **c**. Quantification of total oligodendrocyte lineage cell density in PBS- and LPS-treated aged mice at 7dpl and 14dpl upon lysolecithin-induced spinal cord demyelination (n=5-6 for 7dpl, n=9 for 14dpl, Student’s t-test). **d**. Quantification of the density of proliferating OPCs (OLIG2^+^KI67^+^) in PBS and LPS-treated aged mice at 7dpl and 14dpl upon lysolecithin-induced spinal cord demyelination (n=5-6 for 7dpl, n=8-9 for 14dpl, Student’s t-test). **e.** Representative immunofluorescence images of mature oligodendrocytes labelled with DAPI (white), OLIG2 (green) and ASPA (magenta) in lysolecithin-induced demyelinating lesions in PBS -and LPS-treated mice. Demyelinated area is demarcated by the yellow line. Scale bar=100µm; Inset scale bar=10µm. **f**. Quantification of the density of mature oligodendrocytes (OLIG2^+^ASPA^+^ cells) in PBS and LPS treated aged mice 14 dpl upon lysolecithin-mediated demyelination in PBS and LPS treated mice (n=9-10, Student’s t-test). **g.** Representative immunofluorescence images of myelin-wrapped and damaged axons labelled with DAPI (white), NFH (green), MBP (magenta) and SMI32 (cyan) in PBS and LPS-treated aged mice 14 days after demyelination. Lesion area is drawn in yellow. Scale bar=100µm; Inset scale bar=10µm). **h, i**. Quantification of the density total axons (total NFH^+^) (**h**) and myelin-wrapped axons (MBP^+^NFH^+^)(**i**) in the lesion 14dpl upon lysolecithin-induced demyelination in PBS and LPS treated mice (n=9-10, Student’s t-test). **j-l.** Quantification of the density of total damaged axons (NFH^+^SMI32^+^) (**j**), damaged myelin-wrapped axons (NFH^+^SMI32^+^MBP^+^) (**k**), and damaged unmyelinated axons (NFH^+^SMI32^+^MBP^-^) (**l**) in the lesion 14dpl upon lysolecithin-induced demyelination in PBS- and LPS-treated mice (n=9-10, Student’s t-test).

The extent of demyelination was not affected by LPS pre-exposure (**Sup. Fig 5a**). Microglia response and myelin debris clearance, assessed by immunostaining for IBA1 to identify microglia and macrophages together with an antibody for a degraded epitope of MBP (dMBP)^52^, were also unchanged (**Sup. Fig. 5b-d**). Similarly, LPS-mediated EMI did not alter oligodendrocyte lineage cell density or the density of proliferating OPCs, quantified by co-staining of lineage marker OLIG2 and KI67 (**Fig 6b-d**). Given the well-established delay in OPC differentiation and remyelination in aged rodents, we next focused only on the 14dpl timepoint to evaluate these processes. In contrast to the lack of effect on OPC number and proliferation, we detected a significant increase in the density of differentiated OLIG2^+^ASPA^+^ oligodendrocytes in the demyelinated area of LPS-treated mice at 14dpl (**Fig. 6e, f**). LPS pre-treatment did not alter total axonal density (NFH^+^ axons) or the density of axons surrounded by a ring of myelin (NFH^+^MBP^+^ axons) (**Fig. 6g-i**). However, LPS pre-exposure significantly reduced axonal damage, assessed by immunohistochemistry against dephosphorylated neurofilament (SMI32) in combination with NFH and MBP (**Fig. 6g, j**). This reduction was specific to MBP^+^ myelin-wrapped axons (NFH^+^SMI32^+^MBP^+^), as the density of damaged unmyelinated axons (NFH^+^SMI32^+^MBP^-^) axons was unchanged (**Fig. 6k, l**). These findings suggest that LPS-mediated EMI accelerates OPC differentiation in the aged CNS and promotes the formation of protective myelin that limits axonal damage upon demyelination.

To determine whether this enhanced regenerative response in OPCs required tolerance-like EMI or it is a consequence of persistent chromatin accessibility alterations, we repeated the same paradigm but exposing aged mice to Poly(I:C), which induced long-lasting chromatin accessibility changes but not tolerance-like EMI. Poly(I:C) pre-exposure did not affect the extent of demyelination, microglial response, myelin debris clearance, oligodendrocyte lineage cell density or OPC proliferation at 14dpl (**Sup. Fig. 6a-e**). Unlike LPS, Poly(I:C) pre-exposure did not enhance OPC differentiation into mature oligodendrocytes or reduce axonal damage (**Sup Fig. 6f-i**). Therefore, Poly(I:C) driven enduring changes in chromatin accessibility in the absence of tolerance-like EMI are insufficient to improve aged OPC regenerative capacity.

## Discussion

As long-lived progenitors, OPCs encounter diverse inflammatory signals throughout life, yet how they adapt to inflammation remains unclear. Here, we demonstrate that adult OPCs harbour epigenetic memory of inflammation (EMI) through enduring changes in chromatin accessibility that shape their future inflammatory and regenerative responses.

In response to acute inflammation induced by LPS or Poly(I:C), OPCs rapidly adopt an immune or disease-associated transcriptional programme, described for several neurodegenerative diseases^15,16^. However, these transcriptional changes returned to baseline once inflammation subsided, suggesting that disease-associated states in OPCs are dynamic and tightly coupled to active inflammation^14,24^. But do OPCs, like other long-lived progenitors, harbour a memory of this experience in the form of persistent chromatin remodelling^20–23^?

We demonstrated that adult OPCs undergo long-lasting changes in chromatin accessibility following exposure to LPS and Poly(I:C), even in the absence of ongoing inflammation. Unlike IFNγ-treated neonatal OPCs^24^, LPS pre-exposed adult OPCs displayed blunted transcriptional and epigenetic responses to repeated LPS, whereas Poly(I:C) recall responses remained unchanged. This divergence may stem from the well-established transcriptomic and proteomic differences between neonatal and adult OPCs^53–55^, underscoring the importance of also studying the adult OPC population. Divergence may also arise from the distinct nature of the inflammatory stimuli. In innate immune cells, the nature of the initiating stimulus determines the form of EMI that develops^23,37,38^, consistent with our results in adult OPCs. Although both elicit persistent chromatin accessibility changes, LPS and Poly(I:C) shared only a subset of these memory DARs. These differences may be partly explained by the distinct receptors though which these stimuli act. LPS primarily acts through toll-like-receptor 4 (TLR4), which has been already implicated in tolerance-like EMI in hematopoietic stem cells^23^, whereas Poly(I:C) predominantly activates TLR3^39^. While both receptors are expressed in OPCs, they can engage divergent downstream signalling pathways^56^, potentially leading to non-overlapping DARs. In addition, the broader inflammatory response elicited by LPS and Poly(I:C) in the CNS differs^40^, which may further contribute to the stimulus-specific EMI observed in adult OPCs. Therefore, it is plausible that the diverse inflammatory experiences encountered throughout the OPCs lifespan may imprint distinct EMI states, ultimately shaping how OPCs respond to subsequent stressors.

The tolerance-like EMI induced by LPS provides new insight into the ongoing debate surrounding how OPC immune states influence neuroinflammation and remyelination^8,57^. While several studies have reported that immune-associated OPC profiles impair their regenerative capacity^13,16,24,57^, others propose that disease-associated genes such as *Serpina3n* may support cell survival and prevent apoptosis during early inflammatory stages^14,58^, thereby facilitating remyelination^8^. Although a controlled inflammatory response is required for adequate myelin regeneration, chronic inflammation can hinder this process^3^. This balance is particularly relevant for the ageing CNS, where low-grade chronic inflammation coincides with a progressive decline in remyelination efficiency^3–5,45,46^. However, whether age-related remyelination impairment is linked to the increased expression of immune genes observed in aged OPCs^27,53–55^ remains unknown.

To address whether EMI influences OPC regenerative capacity, we subjected aged mice pre-exposed to LPS- or Poly(I:C) to toxin-induced demyelination. LPS-mediated EMI enhanced OPC differentiation and reduced axonal damage in myelin-wrapped axons within the demyelinated area in aged mice, whereas Poly(I:C) pre-exposure had no detectable effect. These findings suggest that the tolerance-like response induced by LPS but not Poly(I:C), may help restore regenerative potential in aged OPCs. Ageing is also characterized by a progressive epigenetic drift, and aged OPCs exhibit altered expression of chromatin-remodelling enzymes^45,59^. Thus, besides attenuating excessive OPC immune activation, LPS-mediated EMI may also reshape the chromatin landscape of aged OPCs, promoting a chromatin state that supports aged OPC differentiation and regeneration after myelin damage ^54,60–62^.

Acute inflammation induced EMI provides an evolutionary advantage by promoting pathogen clearance, accelerating tissue repair and preserving tissue integrity. However, chronic or repeated inflammatory insults may instead generate maladaptive memory states that sustain and amplify inflammation, thereby worsening disease progression^34,36^. Such maladaptive inflammatory memory has been demonstrated in astrocytes in EAE^33^, in microglia in Alzheimer’s disease models^37^, and has been suggested in OPCs exposed to IFNγ^24^. Recent evidence has shown that adult human OPCs present epigenetic memory of developmental origin as described for other stem cell populations^63^. Whether adult human OPCs also have inflammatory memory and whether such EMI shapes remyelination in humans, however, remains to be addressed. If translatable to the human CNS, EMI in OPCs (and other CNS cell types), may help explain the marked heterogeneity in remyelination capacity observed among patients with MS^64^ as well as the potential influence of the history of prior infections/inflammatory stressors on disease outcome. Our results demonstrate that LPS-mediated EMI enhances OPC differentiation and preserve axonal health in the aged CNS, suggesting that beneficial forms of OPC EMI may be harnessed for therapeutic purposes. However, such approach would require a deeper understanding of how EMI is established, maintained, and recalled, given the potential negative impact of maladaptive inflammatory memory^32,36,51^. Different transcription factors, histone modifications and DNA methylation patterns have been implicated in short- and long-term EMI in other stem and CNS cell types^23,32,33,51,65^. Elucidating the molecular mechanisms governing EMI in OPCs and its persistence over time, will be essential to therapeutically target beneficial or protective OPC EMI while attenuating maladaptive inflammatory memories. This is an especially relevant goal for neurodegenerative and age-related disorders, where inflammatory dysregulation is a central pathological feature^36,51^.

Collectively, our data demonstrate that adult OPCs encode long-lasting epigenetic memories of inflammatory experiences that shape their immune and regenerative responses. These results position EMI as a potential therapeutic target to mitigate age-associated remyelination failure and suggest that the inflammatory experiences stored by OPCs may contribute to inter-individual variability in remyelination efficiency^64,66^.

### Limitations of the study

Although our epigenetic analysis supports cell-intrinsic epigenetic remodelling in OPCs, LPS pre-exposure in our paradigm could also induce EMI in microglia^37,67^. Previous reports have shown that BCG-mediated inflammatory memory in microglia enhances remyelination by promoting myelin debris clearance^51^. However, we observed no changes in myelin debris clearance, suggesting that the enhanced OPC differentiation observed primarily stems from intrinsic OPC EMI. Considering that other CNS cells are known to harbour EMI, including astrocytes, microglia or neurons^33,37,51,68^, the effects of systemic inflammation on these other CNS cells cannot be excluded. Thus, dissecting the relative contribution of each cell type to CNS inflammatory memory will require approaches that enable cell-type specific manipulation of EMI and longitudinal tracking. Moreover, because bulk RNA and ATAC sequencing do not resolve OPC heterogeneity, we are unable to determine whether the tolerance-like EMI observed reflects memory encoded by discrete/specific OPC subsets, the entire OPC population or by distinct OPC populations being recruited after primary and secondary challenges. Although Poly(I:C) induced persistent changes in chromatin accessibility in OPCs, the absence of altered recall responses in our assays suggests that such remodelling may not constitute inflammatory memory in these contexts or alternatively, may only trigger memory-like responses to alternative stressors. Finally, while we observed OPC EMI across the lifespan, the identity, durability and reversibility of these memories, and whether the mechanisms governing EMI in young OPCs are preserved in aged OPCs remain unresolved.

## METHODS

### Animals

All animal work was conducted according to European Union guidelines and with protocols approved by the Bioethics and Biosecurity committee at the Institute of Neurosciences CSIC-UMH as well as from the Institutional Animal Care and Use Committee (*Dirección General de Agricultura, Ganadería y Pesca from Generalitat Valenciana*, protocol 2023-VSC-PEA 0192). All mice were C57BL/6J background from both sexes bred in-house. Mice were kept in a controlled environment with constant temperature (23°C) and humidity (40-60%), on 12h light/dark cycles, with food and water ad libitum. 2-3 months young adult mice were subjected to intraperitoneal injection of PBS, 1mg/kg Lipopolysaccharide (LPS from Escherichia coli O111:B4 Sigma-Aldrich) or 10mg/kg polyinosine-polycytidylic acid (Poly(I:C) high molecular weight, Invivogen). Young adult mice were used for OPC isolation from brain, followed by *in vitro* experiments, RNA sequencing, ATAC sequencing or qPCR. Spinal cords were collected, fixed in 4% PFA for 72h at 4 °C, cryoprotected in 30% sucrose for an additional 72h, embedded in OCT, and sectioned at 12µm for immunohistochemistry.

12-month-old mice were also subjected to intraperitoneal injection of PBS, 1mg/kg LPS or 10mg/kg Poly(I:C) and 5 weeks later underwent spinal cord demyelination induced by lysolecithin injection.

### Lysolecithin-induced spinal cord demyelination

Spinal cord demyelination was induced in the ventral white matter of the spinal cord between vertebrae T11-12 or T12-13, by injecting 1.2 µL of 1% (w/v) L-α-Lysophosphatidylcholine (Lysolecithin; Sigma-Aldrich) under general gaseous anaesthesia. At 7 or 14days post lesion (dpl), mice were anaesthetised with a lethal dose of ketamin-xilacin (Richter pharma, Calier) and transcardially perfused with phosphate buffered saline (PBS) and 4% paraformaldehyde (PFA) (Sigma-Aldrich). Spinal cords were dissected and incubated overnight in 4% PFA at 4 °C, and for 72h in 30% sucrose (Sigma-Aldrich) in PBS before embedding in OCT (Tissue-Tek). Spinal cords were cryosectioned at 12 µm for immunostaining.

### Immunohistochemistry

Spinal cord sections were dried for 30 min at RT and washed twice in TBS-T (Tris-buffered saline with 0.25% Tween20, Sigma-Aldrich). Afterwards, slides were immersed in pre-warmed antigen retrieval solution (1X citrate buffer pH 6.0, Sigma-Aldrich) for 5min at 85°C. After cooling for 30min, slides were washed twice with TBS-T and permeabilized with 1% Triton X-100 in TBS-T for 30 min at RT. Slides were then washed with TBS-T and incubated with blocking solution (5% donkey serum; Sigma-Aldrich) diluted in TBS-T for 1 h at RT in a humidified chamber. Slides were next incubated with primary antibodies overnight at 4 °C and then washed in TBS-T three times. Secondary antibodies and DAPI (1µg/mL; Sigma-Aldrich) were added and incubated for 1h at RT. Finally, slides were washed with TBS-T twice and mounted with fluoromount G. Primary antibodies used: OLIG2 goat (1:500, Biotechne), OLIG2 mouse (1:200, Millipore), PDGFRα rabbit (1:500, Cell Signalling), PDGFRα goat (1:200, Biotechne), ASPA rabbit (1:400, Millipore), SERPINA3N goat (1:500, Biotechne), H3K27ac rabbit (1:500, Abcam), H3K27me3 rabbit (1:500, Cell Signalling), KI67 rat (1:300, eBioscience), NFH chicken (1:1000, Abcam), MBP rat (1:1000, Biorad), SMI32 mouse (1:1000, Biolegend), IBA1 goat (1:500, Abcam) and dMBP rabbit (1:2000 Millipore). Secondary antibodies used: donkey anti-goat AF488, donkey anti-mouse AF555, donkey anti-mouse AF488, donkey anti-rat AF555, donkey anti-goat AF647, donkey anti-rabbit AF555, donkey anti-mouse AF647 (all 1:500; ThermoFisher Scientific) and donkey anti-chicken AF488 (1:500, Jackson ImmunoResearch). Images were taken using Zeiss Axioscan 7 or Zeiss LSM 880 Airyscan Elyra PS1. Further processing was done using Huygens deconvolution software (Huygens Professional version 21.10, Scientific Volume Imaging, The Netherlands, http://svi.nl, RRID: SCR_014237) for H3K27ac and H3K27me3 immunostainings. The analysis was done either manually by two blinded assessors or using Cell Profiler 4.2.1 and Cell Profiler Analyst software. H3K27ac and H3K27me3 distribution analysis of the deconvoluted images was performed as previously described using Fiji Sholl analysis plugin^29,30^. The deconvolved nuclei were maximum projected along the z-axis and transformed to binary (black pixels belonging to any H3K27me3 or H3K27ac signal and the absence of signal of the background belonging to white pixels). Sholl analysis was performed (v4.2.1) on the binary nuclei creating concentric circles from nucleus centre to periphery using a starting radius of 1µm up to 5 µm and 0.035µm radius step size. The number of intersections were plotted in the y axis along the distance from nucleus center and fitted using the 6th degree polynomial function and analyzed using 2-way ANOVA. The area under the curve was calculated for the polynomial function of each nucleus in GraphPad Prism (GraphPad Software, Inc. version 9) and analyzed using 1-way ANOVA.

### OPC isolation and culture

OPCs were isolated from 3–4-month-old mice brain based on PDGFRα expression. Mice were cervically dislocated, and brain was placed in Hibernate-A (ThermoFisher Scientific). Dissected brain was minced using scalpels and transferred to a 15 mL tube with Hanks Buffered Salt Solution (HBSS, Capricorn Scientific) and centrifuged for 1 min at 100 g and 4 °C. The pellet was resuspended in 5 mL dissociation media containing 165U of Papain (Worthington) in hibernate A for 30 min at 37 °C. Papain was washed off with HBSS and cells were centrifuged for 5 min at 300 g and 4 °C. The pellet was then triturated gently with 4 mL trituration buffer (Hibernate-A with 2% B27; ThermoFisher Scientific and 4mM sodium pyruvate; ThermoFisher Scientific) using a 5 mL pipette. After letting the tissue settle for 2 min the supernatant was transferred to a clean 50 mL tube through a 70 µm strainer (Corning). Remaining tissue was triturated gently using a glass polished pipette with another 2 mL of trituration buffer, tissue was left to settle, and supernatant was transferred to the tube through the strainer. This process was repeated another time with a glass fire-polished pipette of decreased diameter and then with a 1 mL pipette. Afterwards, 11mL of 90% Percoll (Sigma-Aldrich) diluted in 10X PBS (ThermoFisher Scientific) was added and topped up to a final volume of 45 mL with Dulbecco’s Modified Eagle’s medium (DMEM) (ThermoFisher Scientific) to remove cell debris. The cell suspension was centrifuged at 800g for 20 min at 4 °C and brake 0. The supernatant was removed, and the cell pellet was washed with HBSS and then resuspended in 10 mL magnetic-activated cell sorting (MACS) buffer (Hibernate-A with 4 mM sodium pyruvate, 2% B27, 0.5% BSA; ThermoFisher Scientific, 2 mM EDTA; Sigma-Aldrich and 10 µg/mL insulin; Sigma-Aldrich). Cell suspension was incubated for 30 min at 37 °C in a 10 cm petri dish to remove microglia. The supernatant was then collected and centrifuged at 300 g for 5 min at 4 °C. The pellet was resuspended in 80µL MACS buffer and incubated with 10µL of FcR blocking reagent (from PDGFRα Microbead Kit, Miltenyi Biotec) for 10 min in ice, then 10µL of PDGFRα microbeads (from PDGFRα Microbead Kit, Miltenyi Biotec) were added and incubated for 15 min in ice. Then the antibody suspension was washed with HBSS and centrifuged at 300 g for 5 min at 4 °C. Finally, cells were resuspended in 500µL MACS buffer and placed in a MACS mini column (Miltenyi Biotec) on a MiniMACS Separator (Miltenyi Biotec) and washed with 1500µL MACS buffer. After the liquid passed through the column, it was placed in a new 1.5ml RNAse-free tube and 1 mL of MACS buffer with 4U/µL RNAsin (Promega) was added to elute the OPCs for RNA and ATAC sequencing experiments.

For *in vitro* experiments, column was eluted using 1mL OPC media containing DMEM, 2% B27, sodium pyruvate (2 mM), insulin (5 µg/mL), Trace Elements B (0.01%; Corning), Forskolin (5 µM; Sigma-Aldrich), Biotin (10 ng/mL; Sigma-Aldrich), Penicillin-streptomycin-glutamine (1%; ThermoFisher Scientific), N-acetyl cysteine (60 µg/mL; Sigma-Aldrich) and 1% SATO stock solution (bovine serum albumin (BSA) fraction V (0.1 mg/mL; ThermoFisher Scientific), sodium selenite (4 μg/mL; Sigma-Aldrich), putrescine (1.61 mg/mL; Sigma-Aldrich), apo-transferrin (0.1 mg/mL; Sigma-Aldrich) and progesterone (4 µg/mL; Sigma-Aldrich)).

For transwell assays, OPCs were plated on OPC media at a density of 20000 cells per well in 8µm transwell insert (CellQART) on 24-well plates (Falcon) with coverslips previously coated with 0.01% Poly-L-ornithine (Sigma-Aldrich) and 10 µg/mL laminin (Sigma-Aldrich) diluted in DMEM. After 16h transwell insert were removed and OPCs in the plate were fixed for 15min with 4% PFA, that was washed with PBS twice before staining with DAPI (Sigma-Aldrich). Images of whole coverslips were acquired using Leica Thunder Imager at 10X magnification and manually quantified.

For primary culture, OPCs were plated on OPC media supplemented with PDGF-AA (20ng/mL, Peprotech) and NT3 (10ng/mL, Peprotech) at a density of 10000 cells per well in 96-well plates (Falcon) previously coated as described. OPCs were maintained with growth factors for 5 days to allow cell recovery with media changes every other, in which 60% of the media was removed and fresh media with PDGF-AA and NT3 added. Six days after growth factor removal (at 11 days after plating) OPCs were fixed for 15min with 4% PFA, that was washed with PBS twice. For immunostaining, cells were blocked with 5% donkey serum (Sigma-Aldrich) and 0.1% Triton X-100 (Sigma-Aldrich) in PBS for 1 h at RT. Primary antibodies were then added diluted in blocking solution and incubated overnight at 4°C. Primary antibodies used were: OLIG2 goat (1:500, Bio-techne), CNPase mouse (1:500, Sigma-Aldrich, clone 11-5B), MBP rat (1:500, Millipore, clone 12) and KI67 rabbit (1:300, Abcam, clone SP6). Afterwards, cells were washed with PBS three times and incubated for 1 h at room temperature with secondary antibodies diluted in blocking solution. Secondary antibodies used included Alexa fluor (AF) 488 donkey anti-rabbit (1:500; ThermoFisher Scientific), AF568 donkey-anti-rat (1:500; Abcam), AF647 donkey anti-mouse (1:500; Abcam) and AF755 donkey anti-goat (1:500; ThermoFisher Scientific) and DAPI (1:10,000; Sigma-Aldrich). Images were acquired using the CellInsight CX5 high content imaging system (ThermoFisher Scientific), with 25 separate fields of view of each well. Cell populations were quantified using CellInsight CX5 analysis software.

### RNA sequencing and RT-qPCR

Isolated OPCs were centrifuged at 1000g for 7min at 4°C to remove media and resuspended in 1 mL of TRI reagent (Sigma-Aldrich), vortexed and incubated for 5 min at room temperature (RT). Then we added 200µL of chloroform (Sigma-Aldrich), vortexed and incubated for 5 min on ice. Samples were centrifuged at 12000g for 15 min at 4°C, and the upper aqueous layer was transferred to a new 1.5mL RNAse-free tube with 2µg Glycogen RNA Grade (ThermoFisher Scientific) and 500µL isopropanol (PanreacAppliChem). After vortexing, the mix was incubated overnight at -20°C and for 10min at RT the next day, then it was centrifuged at 12000g for 10 min at 4°C. Supernatant was discarded, and pellet was resuspended in 1mL of 75% EtOH (Sigma-Aldrich) and centrifuged again at 7000g for 5 min at 4°C. Finally, supernatant was discarded, and RNA pellet was resuspended in 20µL RNAse free H_2_O (ThermoFisher Scientific) and kept at -80°C.

For RT-qPCR, RNA concentration was measured using NanoDrop (ThermoFisher Scientific) before retrotranscription to cDNA with RevertAid Reverse Transcriptase (ThermoFisher Scientific). RT-qPCR was performed using with PyroTaq EvaGreen (Cultek) in a QuantStudio 3 thermocycler (Applied Biosystems). Primers used were: *Tnf* (F:CCTGTAGCCCACGTCGTAG, R:GGGAGTAGACAAGGTACAACCC), *C4b* (F:GGAGAGTGGAACCTGTAGACAG, R:CACTCGAACACGAGTTGGCTTG), *Serpina3N*(F:GCCTCGTCAGGCCAAAAAG, R:TGAACGTGTCAAGAGGGTCAA), *β2M*(F:TCACCCCCACTGAGACTGATA, R:CTTGATCACATGTCTCGATCCCA). *18S* was used as reference gene (F:CGGCTACCACATCCAAGGAA, R:GCTGGAATTACCGCGGCT).

For RNA sequencing RNA integrity was determined by using Bioanalyzer 2100 system (Agilent Technologies). For library preparation messenger RNA was purified from total RNA using poly-T oligo-attached magnetic beads. After fragmentation, the first strand cDNA was synthesized using random hexamer primers followed by the second strand cDNA synthesis. The library was ready after end repair, A-tailing, adapter ligation, size selection, amplification, and purification. The library was checked with Qubit and real-time PCR for quantification and bioanalyzer for size distribution detection. After library quality control, different libraries were pooled based on the effective concentration and targeted data amount, then subjected to Illumina sequencing in a pair-end 150 base pair format and 50 million reads.

### ATAC sequencing

After OPC isolation, 80000 cells were centrifuged at 1000g for 5min at 4°C. Supernatant was removed and pellet was resuspended in 100µL nuclei extraction buffer (NEB) containing 250 mM Sucrose, 25 mM KCl, 5 mM MgCl_2_, 20 mM HEPES-KOH pH 7.8, 0.5% IGEPAL CA-630, 0.2 mM spermine, 0.5 mM spermidine, and 1X protease inhibitors (cOmplete EDTA-free, Roche). Cells were incubated for 10min on ice and then centrifuged at 500g for 7min at 4°C. The pellet was resuspended in NEB without IGEPAL and with 1% BSA, centrifuged again at 1000g for 7min at 4°C and after removing supernatant pellets were resuspended in transposase reaction (TD buffer and Tn5 transposase, Illumina) and incubated at 37 °C for 30 min. Then, DNA was isolated (Qiagen MinElute PCR Purification Kit) and libraries were generated using Diagenode SI tagmented libraries and NEBNext Ultra Q5 Master Mix (New England Biolabs) by PCR. Finally, DNA libraries were purified using AMPure XP beads (Beckman Coulter) to eliminate fragments exceeding 1000bp and primer dimers. To do so, 0.5X of AMPure XP beads were added per sample, incubated for 10min at RT and tubes were placed in a magnetic rack, and after 2 min the supernatant was transferred to a new tube with 1.3X AMPure XP beads. Again, after 10min incubation at RT, tubes were placed in the magnetic rack for 2min and supernatant was discarded. Without moving the tubes from the rack, beads were washed with 200µL of 80% EtOH. EtOH was removed and beads were resuspended in 16µL RNAse free H_2_O, incubated for 10min at RT and placed in the magnetic rack. Supernatant containing DNA libraries was collected and kept at -20°C until sequencing (Novogene).

## Bioinformatic Analysis

### RNA-seq

Raw RNA reads of fastq format were firstly processed through fastp software to obtain clean data by removing reads containing adapter or ploy-N and low-quality reads. All the downstream analyses were based on the clean data with high quality. RNA-seq reads were aligned to the mouse reference genome that was built using Hisat2 (v2.0.5). Raw gene counts were obtained using Rsubread (v 2.20) in paired-end mode with the Mus musculus GRCm38.99 GTF annotation file. Differential expression analysis was performed using DESeq2 (v1.46) and with n=4 individual mice per group (2 females and 2 males). Batch effects were addressed using a likelihood ratio test (LRT) model in DESeq2, and limma (v3.62.2) was used for batch correction in the PCA after batch correction. Gene Ontology enrichment analysis was performed using DAVID Functional Annotation Tools^69^. BEDTools (v2.31.1) and deepTools (v3.5.6) were used to generate bigWig files and heatmaps. Genomic tracks and profiles were visualized using IGV (v2.19.4). The analysis was performed in R 4.4.2 and R.4.5.1. Visualization plots were performed in ggplot2 (4.0.1). GSVA was performed to quantify pathway activity at the sample level. GSVA was run using the GSVA R package (v2.4.4). Enrichment scores were computed using a Gaussian kernel to model continuous, log-transformed expression data. GSVA scores were generated using the default GSVA algorithm without additional smoothing options. GSEA enrichment analysis was performed to assess whether transcriptional programs previous identified a publicly available aged OPC transcriptome^46^ were enriched in our samples. DEGs between young and aged OPCs were extracted aligning the RNA-seq reads to the rat reference genome GRCr8 rat reference genome using Salmon. Differential expression analysis was performed using DESeq2 (v1.46). We created two gene sets with the top 500 DEGs with significantly upregulated and significantly downregulated in aged OPCs vs young OPCs. We performed GSEA using the GSEA Desktop interface for visualization (Broad Institute) (v.4.3.3) with 1000 gene-set permutations, the classic enrichment statistic, and otherwise default GSEA parameters.

### ATAC-seq

ATAC-seq reads were trimmed using trim_galore (v0.6.4_dev) and then aligned to the mouse reference genome mm10 using Bowtie2 (v2.3.5.1). These files were filtered for mapq ≥30 using samtools (v1.1), and duplicate reads were removed using Picard (v2.27). Peak calling was performed using macs2 (v2.1.1.20160306) in BAMPE mode. Differential accessibility analysis was conducted using the R package diffBind (v3.16) which was used to generate a consensus peak set and obtain raw count matrices in all samples. Differential analysis was then performed using DESeq2 and with n=4 individual mice per group for LPS (3 males and 1 female), n=3 individual mice for Poly(I:C) (2 males and 1 female) and the equivalent PBS control mice for each. To correct for batch effects, an LRT model was applied in DESeq2 for DARs, and limma was used for batch correction in the PCA. To identify regions specific to the 5W conditions we used the Elbow package (v1.18.1). A score was calculated by obtaining the lfcSE calculated by Deseq2 in the Wald test and the next formula was used for obtaining the score for the elbow: log2(counts(condition)/counts(PBS))/lfcSE. For annotation of the peaks ChipPeakAnno (v3.24.2) and Biomart (v 2.46.3) were used. To integrate our RNAseq and ATACseq datasets, we quantified the spatial relationship between DARs and DEGs. Continuous DAR-transcription start site (TSS) distances were computed using the distanceToNearest() function, measuring the distance from each DEG to its closest DAR. These distances were used to generate empirical cumulative distribution functions (ECDFs) plots and to assess distributional differences using two-sided Kolmogorov–Smirnov (KS) tests. In parallel, enrichment of DEG-DAR associations was evaluated within five symmetric TSS-centred windows (±1 Kb, ±5 Kb, ±25 Kb, ±50 Kb, ±100 Kb) using hypergeometric test with Benjamini-Hochberg correction for multiple comparisons. Fold enrichment was defined as the ratio of observed to expected DEG-DAR overlaps relative to the expressed gene universe. To account for potential confounding effects of chromosomal gene clustering or accessibility biases, enrichment was further validated using a chromosome aware permutation framework (10 iterations) in which DEG labels were randomly reassigned within each chromosome and empirical permutation scores were calculated. All analyses were implemented in R using GenomicRanges (v1.62.1) and TxDb.Mmusculus.UCSC.mm10.knownGene. To determine potential 3D DAR interactions with DEGs, we integrated our ATAC-seq dataset, with the publicly available H3K27ac HiChIP in OPCs^25^, HiChIP loops linked to the DARs identified between PBS and LPS 5w groups were selected using bedtools pair to bed and these loops integrated with DEGs between LPS 1X and LPS 2X using integrated genomic viewer.

### Statistical analysis

Statistical analyses were performed using GraphPad Prism (GraphPad Software, Inc. version 9) or R. Normality of datasets was tested with Shapiro-Wilk and Kolmogorov-Smirnov tests. Student’s t-test was used to compare between two groups, for example PBS vs LPS in *in vitro* assays. For comparisons between three groups (e. g. PBS, LPS 1X and LPS 5w) one-way analysis of variance (ANOVA) with Sidak’s or Dunnett’s multiple comparison tests was performed comparing always all the groups against PBS. Two-way ANOVA was performed to compare between different treatments and timepoints such as PBS vs LPS at 7 and 14dpl, followed by Sidak’s multiple comparison test. GSVA scores were also compared using two-way ANOVA and Tukey’s multiple comparison test where all the groups were compared against PBS control. For all statistical tests, significant differences were considered at p < 0.05 (*<0.05; **<0.01; ***<0.001; ****<0.0001). Also, in lysolecithin induced-demyelination experiments, animals with lesions smaller than 0.03µm were removed from the analysis, since smaller lesions display a greater regenerative capacity, which may bias the final outcome.

## DATA AVAILABILITY

Bulk RNA-seq and bulk ATAC-seq data of OPCs treated with LPS or Poly(I:C) were deposited at Gene Expression Omnibus (GEO). Raw fastq files, count matrices and genomic tracks are publicly available under accession number GSE324969. Aged OPC transcriptome was downloaded from GSE135765 and the OPC H3K27ac HiChIP was downloaded from GSE166179.

## ACKNOWLEDGEMENTS

We thank the Core Facilities and animal house at the Institute of Neuroscience CSIC-UMH for their technical support, Federico Miozzo for his advice and assistance with ATAC-seq experiments and Francisco J Rodriguez-Baena for his advice and assistance with RNA-seq experiments.

## AUTHOR CONTRIBUTION

The study was conceived by A.G.F. Experiments were designed by A.G.F and S.C.F. Experiments were performed and/or analysed by S.C.F, S.N., A.A.G, J.P.L., A.C.B and J.A.G.S. J.A.G.S., H.C., and A.B. provided advice on experimental design and interpretation. Manuscript was written by S.C.F and A.G.F with contributions from all authors. A.G.F. obtained the funding and oversaw the study.

## DECLARATION OF THE USE OF GENERATIVE AI AND AI-ASSISTED TECHNOLOGIES DURING THE WRITING PROCESS

During the preparation of the manuscript the authors used Microsoft Copilot to revise English grammar and usage. After using this tool, the authors reviewed and edited the content and take full responsibility for the content of the publication.

---

**Supplementary Figure 1:**
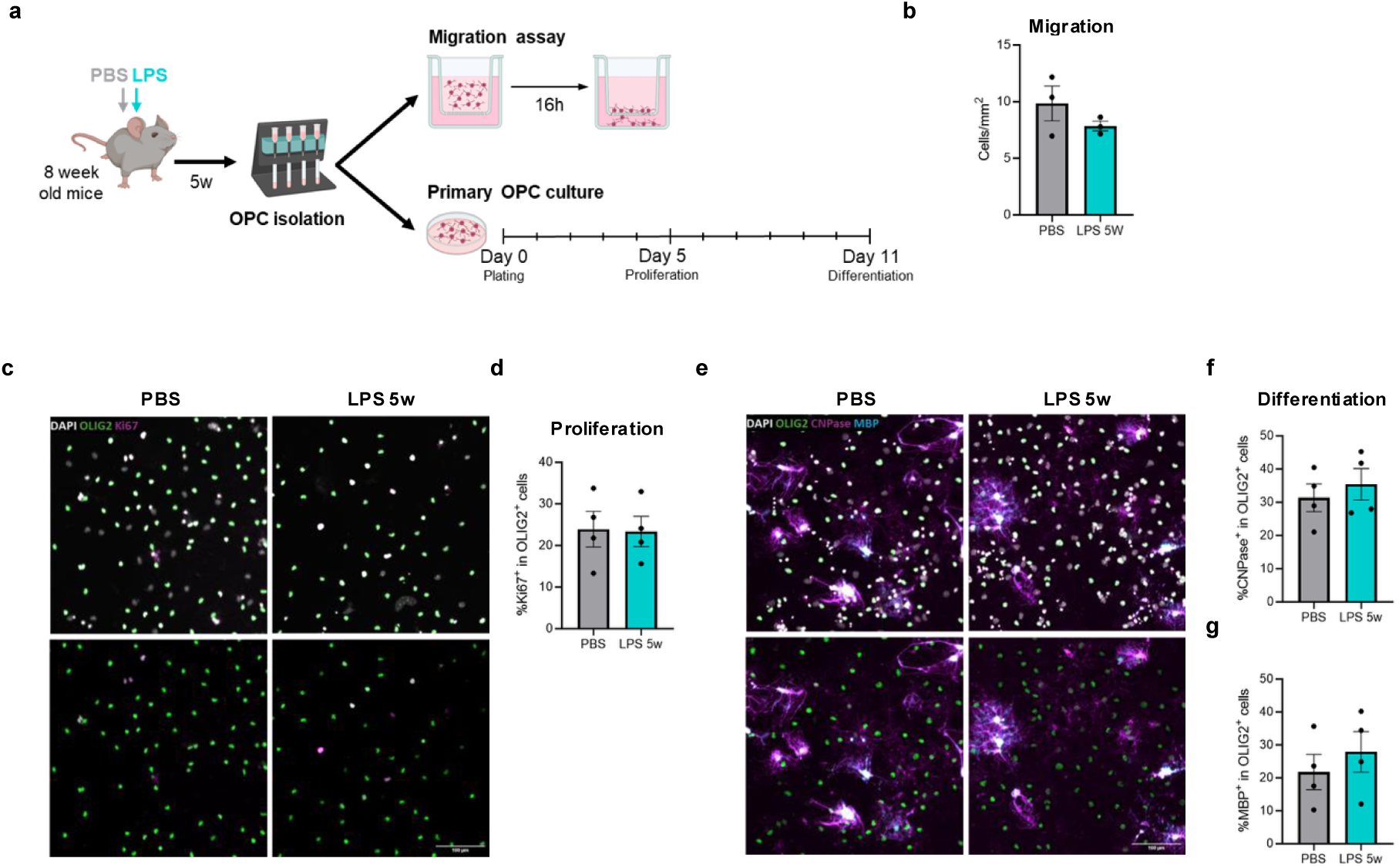
LPS does not modify OPC behaviour *in vitro* 5 weeks after injection. **a.** Schematic diagram of the experimental design to assess OPC migration, proliferation and differentiation capacity 5 weeks after PBS or LPS treatment *in vitro*. **b.** Quantification of OPC migratory capacity indicated as the density of cells per mm^2^ that have migrated through the transwell 16h after isolation and attached to the coverslip below. (n=3 mice, Student’s t-test). **c.** Representative immunocytochemistry images of OPCs isolated 5 weeks after PBS and LPS treatment and cultured *in vitro* labelled with DAPI (white), OLIG2 (green), and KI67 (magenta); scale bar=100µm. **d.** Quantification of the *in vitro* proliferative capacity of OPCs isolated 5-weeks after PBS or LPS treatment indicated as percentage of OLIG2^+^ cell expressing KI67 (n=4 mice, U-Mann Whitney test). **e.** Representative immunofluorescence images of OPCs isolated 5 weeks after PBS and LPS treatment and cultured *in vitro* labelled with DAPI (white), OLIG2 (green), CNPase (magenta) and MBP (cyan); scale bar=100µm**. f, g.** Quantification of the *in vitro* differentiation capacity of OPCs isolated 5-weeks after PBS or LPS, quantified as the percentage of OLIG2^+^ cells expressing CNPase (**f**) or MBP (**g**) (n=4, U-Mann Whitney test).

**Supplementary Figure 2:**
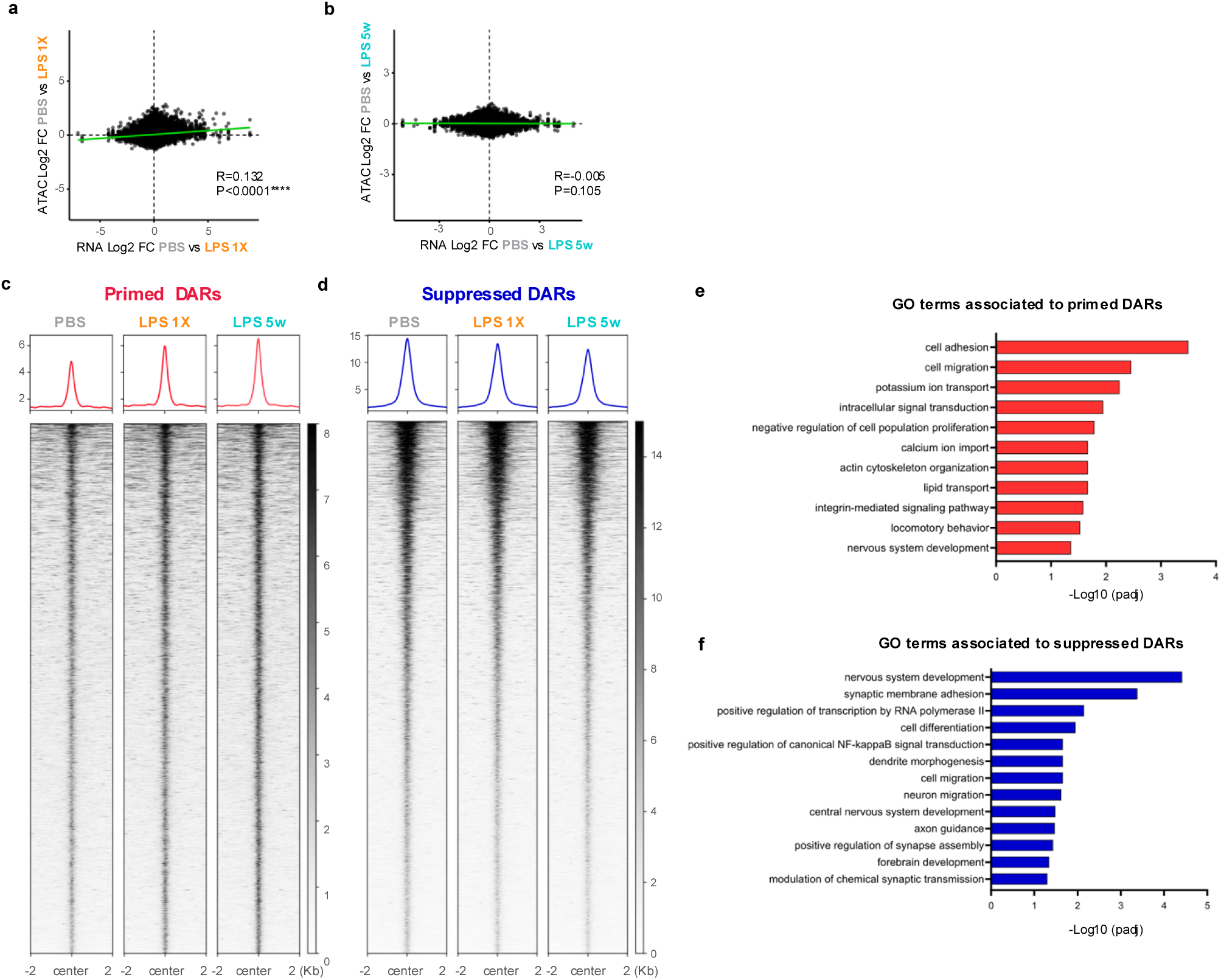
Primed and suppressed memory DARs present similar behaviour but are associated with different GO biological processes. **a, b.** Correlation of the log_2_ fold change in RNA expression and chromatin accessibility between PBS and LPS 1X (**a**) and PBS compared to LPS 5w (**b**). **c.** Heatmap and chromatin accessibility profile showing mean ATAC-seq signal intensity of primed DARs in PBS, LPS 1X and LPS 5w. **d.** Heatmap and chromatin accessibility profile showing mean ATAC-seq signal intensity of suppressed DARs in PBS, LPS 1X and LPS 5w. **e, f**. Bar graph depicting GO biological processes enriched within the genes associated by proximity to primed (**e**) and suppressed (**f**) chromatin regions identified between PBS and LPS 5w.

**Sup Fig3:**
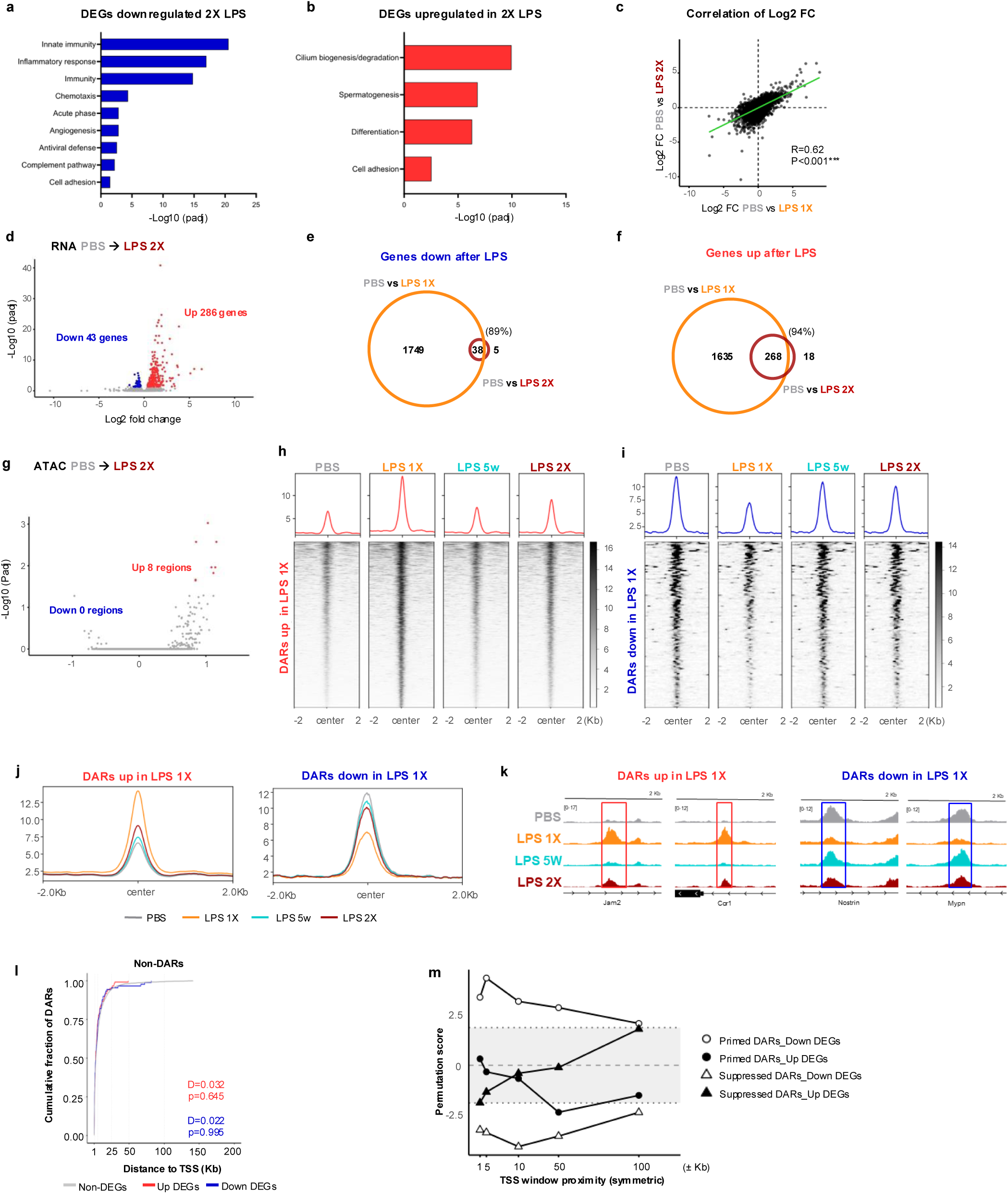
LPS 2X induces a blunted inflammatory response at transcriptional and epigenetic level. **a, b.** GO biological processes significantly enriched amongst the genes downregulated (**a**) and upregulated (**b**) between LPS 2X and LPS 1X OPCs. **c.** Correlation of the log_2_ fold change in expression observed in all the genes shared between PBS and LPS 1X and PBS and LPS 2X. **d.** Volcano plot depicting DEGs identified when comparing OPC transcriptome in PBS versus 2X-LPS (upregulated in red, downregulated in blue). **e, f.** Venn diagrams showing the downregulated **(e)** and upregulated **(f)** genes shared between LPS 2X and LPS 1X when compared to PBS control. **g.** Volcano plot depicting DARs identified between PBS and LPS 2X. **h, i.** Chromatin accessibility profile and heatmap of the mean ATAC signal of the DARs with increased (**h**) and decreased (**i**) accessibility between PBS and LPS 1X showed in PBS, LPS 1X, LPS 5w and LPS 2X. **j, k.** Chromatin accessibility profile showing the log_2_ chromatin accessibility changes of the DARs that show an increased (**j**) or decreased (**k**) chromatin accessibility in LPS 1X compared to PBS. The profile depicted the mean chromatin accessibility in OPCs from the PBS, LPS 1X, LPS 5w and LPS 2X. **l.** Representative IGV profile images of the merged ATAC signal of example genomic loci identified as DARs between PBS and LPS 1X. Merged ATAC-seq profiles from four independent samples in PBS, LPS 5w, LPS 1X and LPS 2X conditions are shown. Genes associated with these chromatin regions are as indicated. **m.** Empirical cumulative distribution function (ECDF) plots showing the cumulative fraction of genes as a function of their distance to the nearest non-DAR peak. Curves for DEGs and non-significant genes are compared visually and tested using the Kolmogorov–Smirnov test. **n.** Permutation score trajectories illustrating the window-sensitivity of DAR–DEG proximity enrichment. Observed DAR–DEG overlaps are compared to empirical null distributions across symmetric TSS windows; the grey band denotes the non-significant region (|σ| ≤ 1.96), with dotted lines marking the permutation-based significance threshold.

**Supplementary Figure 4:**
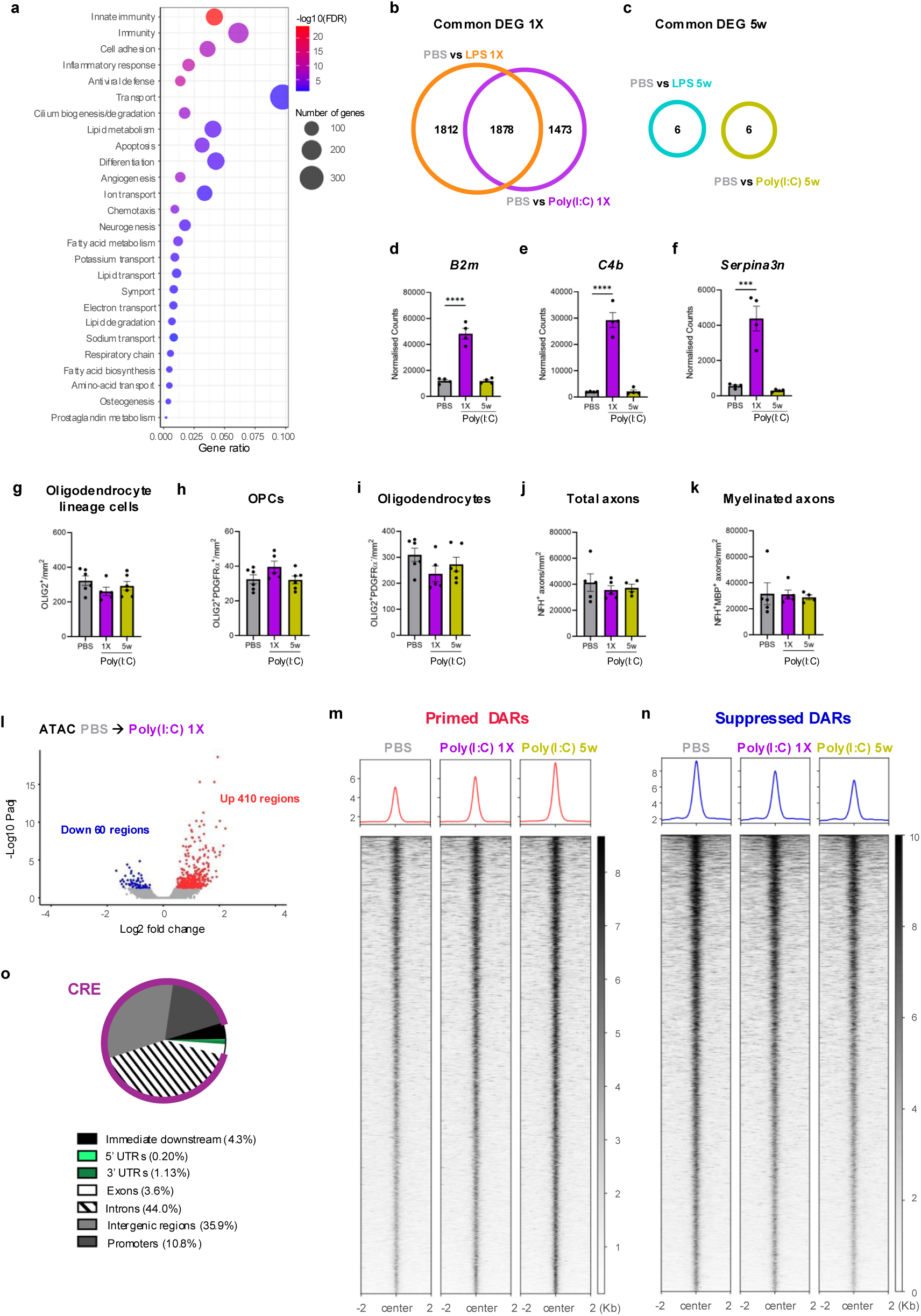
OPCs display stimulus-specific transcriptomic and chromatin alterations following Poly I:C or LPS exposure. **a.** Bubble plot depicting the GO biological processes significantly enriched among DEGs in OPCs 24h after Poly(I:C) treatment. **b, c.** Venn diagram showing the DEGs shared between LPS- and Poly(I:C)-induced transcriptomic OPC response 24h **(b)** and 5w **(c)** after the stimulus. **d-f.** Bar graphs showing the normalised counts of disease-associated OPC markers (*B2m, C4b, Serpina3n*) in PBS, Poly(I:C) 1X and Poly(I:C) 5w (n=4, 1way-ANOVA, Dunnet’s posthoc test). **g-I.** Quantification of the density of total oligodendrocyte lineage cells (OLIG2^+^) (**g**), OPCs (OLIG2^+^PDGFRα^+^) (**h**), and mature oligodendrocytes (OLIG2^+^PDGFRα^-^cells) (**i**) in the spinal cord white matter in PBS, Poly(I:C) 1X and Poly(I:C) 5w mice (n=6 mice, 1-way ANOVA, Dunnet’s posthoc test). **j, k.** Quantification of the density of total axons (NFH^+^) (**j**) and myelin wrapped-axons (NFH^+,^ MBP^+^) (**k**), in the spinal cord white matter in PBS, Poly(I:C) 1X and Poly(I:C) 5w mice (n=6, 1-way ANOVA, Dunnet’s posthoc test). **l.** Volcano plot showing DARs in Poly(I:C) 1X when compared to PBS. DARs with increased chromatin accessibility are shown in red while DARs with diminished chromatin accessibility are depicted in blue. **m,n.** Heatmap and chromatin accessibility profile showing ATAC-seq signal intensity of primed (**m**) and suppressed (**n**) DARs in PBS, Poly(I:C) 1X and Poly(I:C) 5w. **o.** Pie chart indicating the type of DNA regions identified as memory DARs (primed and suppressed) in the Elbow plot comparing Poly(I:C) 5w and PBS.

**Supplementary Figure 5:**
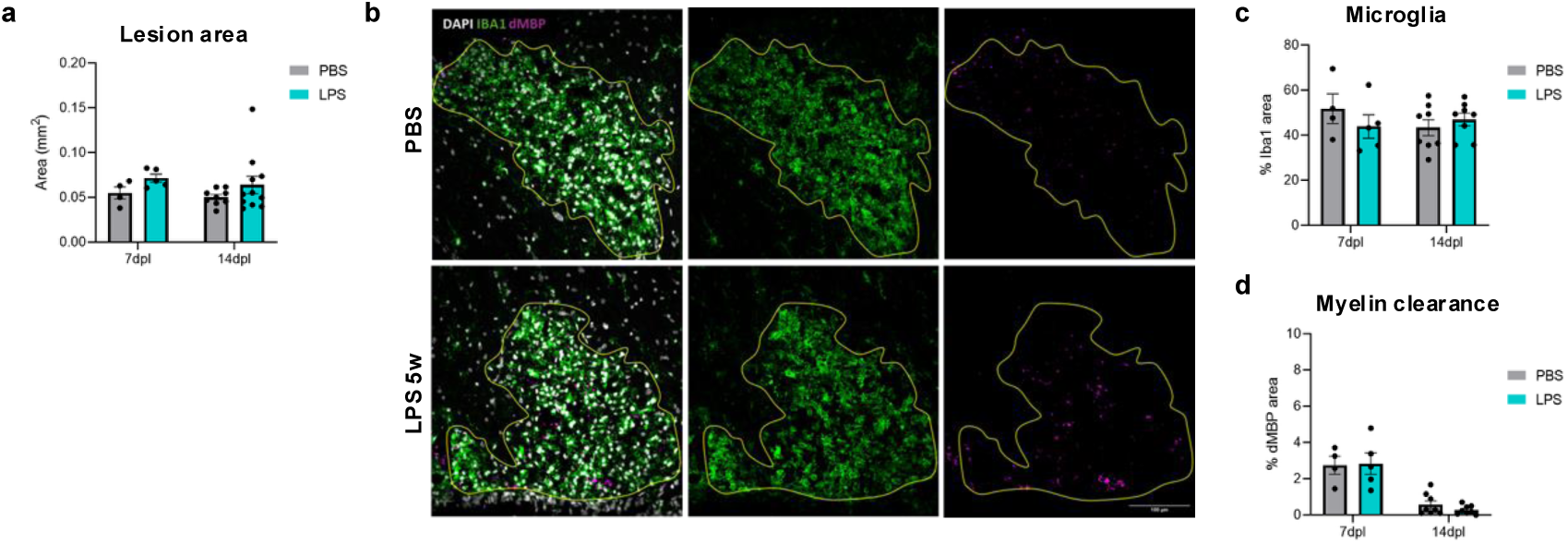
LPS pre-exposure does not affect myelin debris clearance upon demyelination. **a.** Quantification of the demyelinated lesion area in PBS and LPS-treated aged mice at 7 and 14 days post-lesion (n=4-5 at 7dpl and n=9-10 at 14dpl, 2-way ANOVA, Dunnet’s posthoc test). **b.** Representative immunofluorescence images of microglia/macrophage responses and myelin debris clearance labelled with DAPI (white), IBA1 (green) and dMBP (magenta) in PBS and LPS-treated aged mice 7 and 14 dpl upon lysolecithin-induced demyelination. Lesion area is drawn in yellow. Scale bar=100µm). **c, d**. Quantification of IBA1^+^ cell density (**c**) and myelin debris clearance (dMBP) (**d**) in PBS- and LPS-treated aged mice 7 and 14 dpl after lysolecithin-induced spinal cord demyelination (n=4-5 at 7dpl and n=9-10 at 14dpl, 2-way ANOVA, Dunnet’s posthoc test).

**Supplementary Figure 6:**
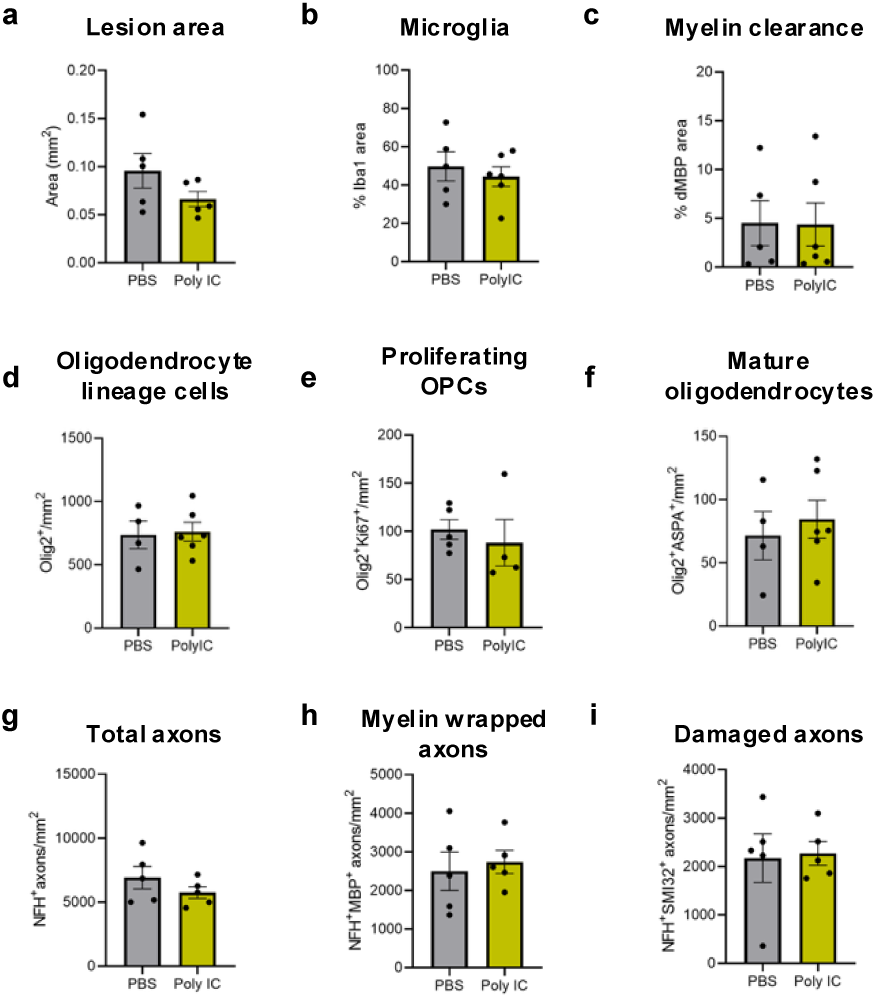
Poly(I:C)-mediated persistent chromatin accessibility changes do not modify OPC response to lysolecithin induced demyelination. **a-c.** Quantification of the demyelinated lesion area (**a**), the percentage of lesion area occupied by IBA1^+^ microglial/macrophage cells (**b**) and the percentage of lesion area occupied by dMBP as a proxy for myelin debris clearance (**c**) 14dpl upon lysolecithin-induced demyelination in PBS- and Poly(I:C)- treated aged mice (n=5-6, U-Mann Whitney). **d-f.** Quantification of the density of total oligodendrocyte lineage cells (**d**), proliferating OPCs (**e**) and mature oligodendrocytes (**f**) 14 dpl upon lysolecithin-induced demyelination in PBS- and Poly(I:C)- treated aged mice (n=5-6, U-Mann Whitney). **g-h.** Quantification of total axons (**g**), myelin wrapped axons (**h**) and total damaged axons **(I)** in PBS and Poly(I:C)-treated aged mice at 14 days post-lesion (n=5-6, Student’s t-test).

